# Nucleosome patterns in four plant pathogenic fungi with contrasted genome structures

**DOI:** 10.1101/2021.04.16.439968

**Authors:** Clairet Colin, Lapalu Nicolas, Simon Adeline, Jessica L. Soyer, Viaud Muriel, Zehraoui Enric, Dalmais Berengère, Fudal Isabelle, Ponts Nadia

## Abstract

Fungal pathogens represent a serious threat towards agriculture, health, and environment. Control of fungal diseases on crops necessitates a global understanding of fungal pathogenicity determinants and their expression during infection. Genomes of phytopathogenic fungi are often compartmentalized: the core genome contains housekeeping genes whereas the fast-evolving genome mainly contains transposable elements and species-specific genes. In this study, we analysed nucleosome landscapes of four phytopathogenic fungi with contrasted genome organizations to describe and compare nucleosome repartition patterns in relation with genome structure and gene expression level. We combined MNase-seq and RNA-seq analyses to concomitantly map nucleosome-rich and transcriptionally active regions during fungal growth in axenic culture; we developed the MNase-seq Tool Suite (MSTS) to analyse and visualise data obtained from MNase-seq experiments in combination with other genomic data and notably RNA-seq expression data. We observed different characteristics of nucleosome profiles between species, as well as between genomic regions within the same species. We further linked nucleosome repartition and gene expression. Our findings support that nucleosome positioning and occupancies are subjected to evolution, in relation with underlying genome sequence modifications. Understanding genomic organization and its role in expression regulation is the next gear to understand complex cellular mechanisms and their evolution.

## Introduction

Fungi account for a huge part of the Earth biodiversity with a current estimate of 2.2 to 3.8 million species (Hawksworth and Lücking, 2017). Fungi are organisms of major environmental importance as they develop beneficial symbiotic associations with plants and are able to decay dead organic matter (Taylor and Osborn, 1996). Unfortunately, fungi are also very efficient pathogens causing important damages in agriculture, human health, and the environment (Fisher *et al*., 2012). Control of fungal diseases on crops necessitates a global understanding of fungal pathogenicity determinants and the control of their expression during infection. Among these pathogenicity determinants, fungi secrete an arsenal of molecules known as effectors, key elements of pathogenesis which modulate innate immunity of the plant and facilitate infection. Effectors can be small proteins, secondary metabolites and small RNAs (Weiberg *et al*., 2013; Lo Presti *et al*., 2015; Collemare, O’Connell and Lebrun, 2019). Upon plant infection, fungi undergo a tightly controlled transcriptional reprogramming, and different sets of effectors are expressed at specific stages of pathogen development and host colonization (Toruño, Stergiopoulos and Coaker, 2016; Zhao *et al*., 2021; Ding, Gardiner and Kazan, 2022). Plant-associated fungi generally show contrasted genomic landscapes including ‘plastic’ *loci* with a high prevalence of transposable elements (TE). These genomes either show an overall large proportion of TE evenly distributed throughout the genome, or TE clustered in specific regions such as long TE-rich blocks, accessory chromosomes or subtelomeric areas (Sánchez-Vallet *et al*., 2018). Effector genes are over-represented in these TE-rich regions. TE-rich compartments have heterochromatin properties contrary to TE-poor regions which have euchromatin properties. The location of effector genes in regions enriched in TEs has been shown to provide a tight control of their expression through chromatin remodeling. Indeed, several recent studies pointed out the potential role of chromatin remodeling in the regulation of effector-encoding genes and the control of secondary metabolism (reviewed in Soyer, Rouxel and Fudal, 2015; Collemare and Seidl, 2019).

Eukaryotic chromatin is packaged into nucleosomes, each composed of DNA wrapped around a histone octamer associated with various other proteins, and separated by linker DNA (Richmond and Davey, 2003). These histone proteins are composed of histone core where the DNA is wrapped and histone tails which can be chemically modified by specific enzymes changing the chromatin 3D-structure and DNA accessibility to polymerases and transcription factors (TF). Nucleosome assembly is further stabilized by the binding of a linker histone H1. Positioning of nucleosomes throughout the genome and post-translational modifications of histones have a significant regulatory function by modifying availability of binding sites to TF and to polymerases, affecting DNA-dependent processes such as transcription, DNA repair, replication and recombination (Radman-Livaja and Rando, 2010; Struhl and Segal, 2013). Nucleosome positioning (*i*.*e*., the position of the nucleosome along the DNA sequence) and occupancy (*i*.*e*., a measure of the actual level of occupation of a given position by a nucleosome in a pool of cells) are determined by a combination of DNA sequence features, TF, chromatin remodelers and histones modifiers (see (Singh and Mueller-Planitz, 2021) for a review). Genome-wide maps of nucleosome occupancy and positioning are still sparse in fungi and have only been developed in a few Hemiascomycota yeast species, including *Saccharomyces cerevisiae* (Yuan *et al*., 2005; Tsankov *et al*., 2010), in the ascomycete *Aspergillus fumigatus* (Nishida *et al*., 2009) and the basidiomycete *Mixia osmundae* (Nishida *et al*., 2012). The studies revealed that promoter, enhancer and terminator regions were depleted in nucleosomes, allowing access to TF, and that the nucleosomal DNA length distribution was similar in *M. osmundae* and *A. fumigatus* but differed from that of hemiascomycetous yeasts. No comparative genome-wide analyses of nucleosome positioning have been performed in ascomycetes and notably not in plant pathogenic fungi.

In the present study, we investigated genome-wide nucleosome localization in four different plant pathogenic ascomycetes showing different genomic organizations: i) *Leptosphaeria maculans* ‘brassicae’ (Lmb), a hemibiotrophic pathogen of *Brassica* species, including oilseed rape; ii) the most closely related species of Lmb, *Leptosphaeria maculans* ‘lepidii’ (Lml), a pathogen of *Lepidium* spp.; iii) *Fusarium graminearum*, a hemibiotrophic pathogen of cereals and iv) *Botrytis cinerea*, a polyphagous necrotrophic pathogen causing grey mould on more than 1,400 plant species. The genome of Lmb has been invaded by TE (which represent more than 30 % of its genome) and is composed of alternating compartments: gene-rich GC-equilibrated and TE-rich AT-rich genomic regions (Rouxel *et al*., 2011; Dutreux *et al*., 2018). In contrast, the Lml genome presents only 4 % of repeats which are evenly distributed throughout the genome (Grandaubert *et al*., 2014). Genomes of *F. graminearum* and *B. cinerea* have a very low to low TE-content (Cuomo *et al*., 2007; Amselem *et al*., 2011; King *et al*., 2015; King, Urban and Hammond-Kosack, 2017). The genome of the reference strain of *B. cinerea*, B05.10, contains 4 % of TE, which are localized essentially in the telomeric and centromeric regions of the core chromosomes, or on the two dispensable chromosomes (Amselem *et al*., 2011; Porquier *et al*., 2016), *i*.*e*., chromosomes not essential to immediate survival and missing in some or most individuals (Soyer *et al*., 2018). The genome of *F. graminearum* contains very little TE identified to date (0.3 % (Cuomo *et al*., 2007; King *et al*., 2015; King, Urban and Hammond-Kosack, 2017)).

In this study, we compare nucleosome repartition patterns in relation with genome structure and gene expression level in these four phytopathogenic Ascomycota. To gain insight into the role of nucleosome positioning and occupancy in regulating fungal pathogen transcription, we applied micrococcal nuclease digestion of mono-nucleosomes couple with high-throughput sequencing (MAINE-seq or MNase-seq) with regards to mRNA abundance to concomitantly map nucleosome-rich regions and transcriptionally active regions during fungal growth.

## Materials and Methods

### Strains and culture conditions

The studied fungi were cultured independently in the media and conditions classically used for each of them. *Leptosphaeria maculans* ‘brassicae’ v23.1.3 and *Leptosphaeria maculans* ‘lepidii’ IBCN84 mycelia were inoculated into 100 mL of Fries liquid medium (1 g/L NH_4_NO_3_, 5 g/L C_4_H_12_N_2_O_6_, 1 g/L KH_2_PO_4_, 0.5 mg/L MgSO_4_ 7H_2_O, 130 mg/L CaCl_2_, 100 mg/L NaCl, 30 g/L C_12_H_22_O_11_ and 5 g/L Yeast extract). Tissues were harvested after growing for seven days in the dark at 25°C. *Botrytis cinerea* strain B05.10 (10^6^ spores/mL) was grown for two days on solid Malt Medium (MM, 20 g/L malt extract, 5 g/L yeast extract and 15 g/L agar) covered with a cellophane layer (Simon *et al*., 2013; Kelloniemi *et al*., 2015). The plates were incubated in a growth chamber (Sanyo MLR-350H) at 23°C with an alternation of 14 h of white light and 10 h of darkness. After two days of culture, mycelia were ground in liquid nitrogen and stored to -80°C until further processing. *Fusarium graminearum* strain CBS185.32 (Centraal Bureau voor Schimmelcultures, Utrecht, the Netherlands) was grown for three days in modified liquid MS (glucose was substituted with sucrose) as previously described (Boutigny *et al*., 2009). All cultures were done in three biological replicates.

### Preparation of nucleosomal DNA

Fungal material was harvested and treated with microccocal nuclease (MNase, cat. #MS0247S, New England BioLabs). For Lmb and Lml, ∼300 mg of mycelium were digested with 5 µL of MNase for 10 min at 37°C (Soyer *et al*., 2015), directly followed by DNA purification as previously described (Soyer *et al*., 2020). For *F. graminearum*, mycelia were harvested by filtering and immediately homogenized for 1 min at 30 Hz using a TissueLyzer (Qiagen). Then, 100 mg of ground mycelium was digested for 10 min at 37°C with 15 µL of MNase in 600 µL of digestion buffer (0.6% v/v IGEPAL, 50 mM NaCl, 2 mM Pefabloc, 50 mM Tris-HCl pH8, 10 mM CaCl_2_). The reactions were stopped with 10mM EDTA and the samples treated with RNAse followed by proteinase K prior DNA purification with phenol/chloroform and ethanol precipitation. For *B. cinerea*, we digested 100-200 mg of mycelium per sample with 1 µL MNase at 37°C (Soyer *et al*., 2015) for 10 min. The reactions were stopped by adding 10mM EDTA and samples treated with RNAse A followed by proteinase K. DNA purification was realized with the “Nucleospin Gel and PCR clean up kit” (Macherey Nagel, cat #740609.250). For all samples, nucleic acid quantification was performed by UV spectrometry using a Nanodrop-ND 1000 apparatus, and digestion profiles were checked by 2% agarose gel electrophoresis. Nucleosomal DNA was stored at -20°C until DNA library preparation.

### Extraction of total RNA

For Lmb and Lml, total RNA was extracted from mycelium grown for one week in Fries liquid medium as previously described (Fudal *et al*., 2007). For *F. graminearum*, mycelia were harvested by filtering, rinsed twice with sterile deionized water, and flash frozen in liquid nitrogen. One milliliter of TRIzol_TM_ Reagent (Thermo Fischer Scientific) was added to 200 mg of mycelium before grinding for 1.5 min at 30 Hz using a TissueLyzer (Qiagen). Total RNA was then extracted using a previously published protocol (Hallen *et al*., 2007). For *B. cinerea*, total RNA was extracted from frozen ground mycelium using a previously published protocol (Kelloniemi *et al*., 2015). All total RNA samples were stored at -80°C until preparation of RNA library.

### Preparation of sequencing libraries, high-throughput sequencing, and read pre-processing

MNase-seq libraries were prepared from purified nucleosomal DNA using the kit NEBNext Ultra DNA Library Prep Kit for Illumina (cat. # E7370L New England BioLabs) following the manufacturer’s instructions. The NEBNext Ultra Directional RNA Library Prep Kit for Illumina (cat. # E7420L New England BioLabs) was used to prepare all RNA-seq libraries, following the manufacturer’s instructions. Sequencing was performed by the GenomEast platform, a member of the ‘France Génomique’ consortium (ANR-10-INBS-0009). Samples were run in 9-plex on an Illumina HiSeq 4000 in paired mode, 2×50 bp. Initial read quality check was performed using FastQC (https://www.bioinformatics.babraham.ac.uk/projects/fastqc/). Raw reads were then pre-processed with Trimmomatic v0.32 (Bolger, Lohse and Usadel, 2014) to clip out any remaining sequencing adapter sequence and crop low quality nucleotides (minimum accepted Phred score of 30). Reads in pairs of 40 bp or more in length were used in the present analysis.

### Transcriptome analyses

RNA-seq reads were mapped against their respective reference genomes (see Table 1) using STAR v2.5.1 (Dobin *et al*., 2013). TPM counts (Transcripts Per Million reads, (Li *et al*., 2010)) were computed using the count TPM tool provided with the MNase-Seq Tool Suite (MSTS; Supplementary Figure 1) that was developed in-house to analyse genome-wide nucleosome positioning data combined with RNA-seq data (https://github.com/nlapalu/MSTS).

**Table 1:**
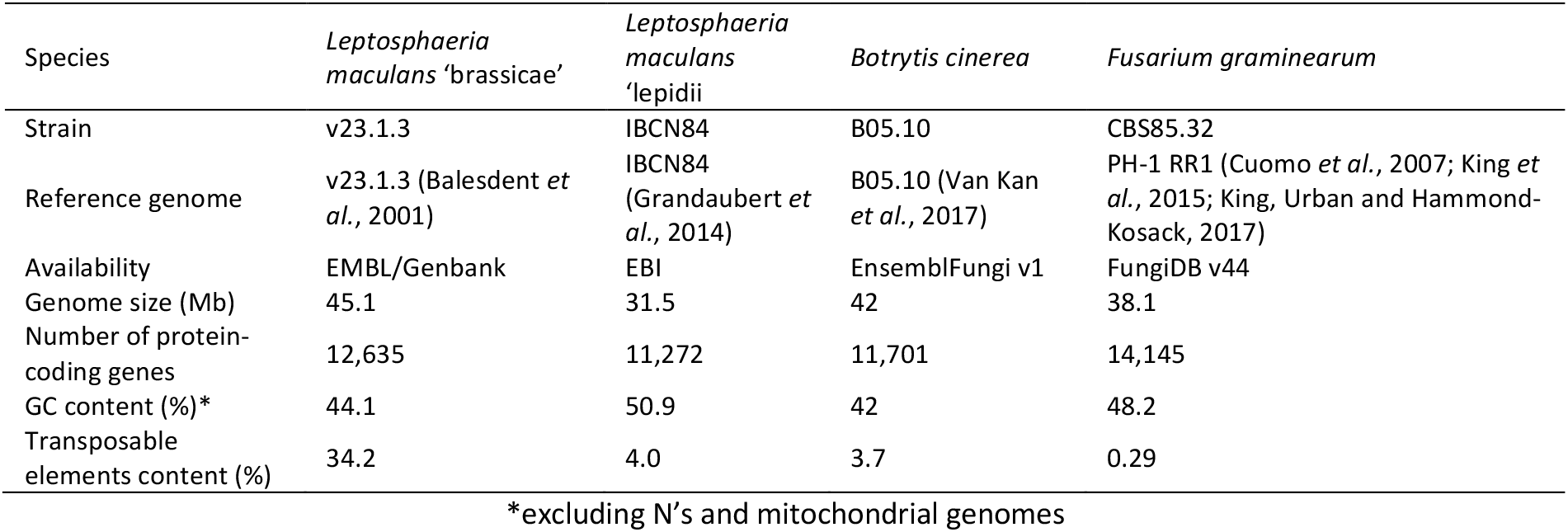
Characteristics of reference genomes for the four fungal species studied.

| Species | *Leptosphaeria maculans* 'brassicae' | *Leptosphaeria maculans* 'lepidii' | *Botrytis cinerea* | *Fusarium graminearum* |
| --- | --- | --- | --- | --- |
| Strain | v23.1.3 | IBC84 | B05.10 | CBS85.32 |
| Reference genome | v23.1.3 (Balesdent *et al.*, 2001) | IBC84 (Grandaubert *et al.*, 2014) | B05.10 (Van Kan *et al.*, 2017) | PH-1 RR1 (Cuomo *et al.*, 2007; King *et al.*, 2015; King, Urban and Hammond-Kosack, 2017) |
| Availability | EMBL/Genbank | EBI | EnsemblFungi v1 | FungiDB v44 |
| Genome size (Mb) | 45.1 | 31.5 | 42 | 38.1 |
| Number of protein-coding genes | 12,635 | 11,272 | 11,701 | 14,145 |
| GC content (%)* | 44.1 | 50.9 | 42 | 48.2 |
| Transposable elements content (%) | 34.2 | 4.0 | 3.7 | 0.29 |

\*excluding N's and mitochondrial genomes

### MNase-seq analyses

MNase-seq paired-end reads were mapped using Bowtie2 software ran in very-sensitive mode (Langmead and Salzberg, 2012). MSTS (MNase-seq Tool Suite) was used to compute all phasograms, dinucleotide composition, as well as nucleosome density profiles of genomic compartments and/or gene lists (Supplementary Figure 1). Lists of near-universal single copy orthologs were obtained by running BUSCO3 for Fungi (Simão *et al*., 2015; Waterhouse *et al*., 2018) on each reference genome studied. Graphical visualisations were computed with MSTS and Matlab R2020b (MathWorks). Frequency distributions of read coverage per base, obtained with the *Phasogram* function of MSTS were scaled (z-score) and plotted with Matlab. For each replicate, phases, standard errors (se), R^2^ (coefficient of determination), and *p*-values (*F*-test) were determined after linear regression fitting to the first four successive peak positions.

## Results and Discussion

### Establishing nucleosome landscapes of the Pezizomycotina *L. maculans* ‘brassicae’, *L. maculans* ‘lepidii’, *B. cinerea*, and *F. graminearum*

We investigated the nucleosome landscapes of four fungal species of the Ascomycota subdivision Pezizomycotina (*L. maculans* ‘brassicae’, *L. maculans* ‘lepidii’, *B. cinerea*, and *F. graminearum*) by MNase-seq (see Supplementary Table 1 for descriptive sequencing metrics). Each experiment was performed using three biological replicates that were sequenced independently at more than 70-fold coverage depths by 147 bp-long nucleosome footprints (defined as the core coverage depths of sequence sufficient for in-depth characterization of nucleosome positioning (Valouev *et al*., 2011)). In order to explore and visualise NGS data obtained from MNase-seq experiments, we developed a collection of utility tools, called MSTS for “MNase-Seq Tool Suite”, assembled in a workflow aiming at profiling nucleosome landscapes in relation to genomic features as well as gene expression (Supplementary Figure 1).

Several tools were previously developed to explore and analyse MNase-seq data such as DANPOS (Chen *et al*., 2013), nucleR (Flores and Orozco, 2011) or CAM (Hu *et al*., 2017). Among the full list of available tools maintained at https://generegulation.org, several tools do not handle paired-end data, were developed for ATAC-Seq, or do not provide visualization, that led us to implement previously published methods in the Python package MSTS. The main features of MSTS are establishment of nucleosome map with nucleosome categorization, comparison with annotation features, Phasogram correlated with gene expression levels or dinucleotides pattern analysis. All tools export results in broad range of graphics and their associated raw data allowing post-process combining several experiments by scaling such as z-score. MSTS is able to consider gene density of small eukaryote genomes like fungal phytopathogens, limiting analysis of phasograms to specific annotation features and avoiding analysis of bases collapsing with other annotated features. This is particularly interesting for NFR analysis at Transcription Start Site (TSS), where the signal could be biased due to the short distance between genes or the overlap of UTRs between adjacent genes. MSTS workflow was applied to our datasets, beginning with the exploration of nucleosome distribution at the genome scale.

### Genome-wide nucleosome spacing

We first explored nucleosome landscapes in the four fungal genomes by measuring the average distance between nucleosomes genome-wide; we computed phasograms, *i*.*e*., frequency distributions of coverage per base genome-wide for all four species (Figure 1 and Supplementary Table 2). Phasograms obtained in nucleosome mapping resemble oscillating sine wave signals, for which period is the length of DNA bound to the histone octamer plus the length of the DNA stretch to the next nucleosome, averaged genome wide (Valouev *et al*., 2011). Phasing signals were observed genome-wide over 1,200 bp sliding windows revealing six to seven nucleosome peaks in a wave signal decaying in intensity with increasing distance and significant linear regression on peak apex positions, as previously described (Valouev *et al*., 2011). We found that, in the fungi studied, nucleosomes are 161 to 172 bp distant from each other (centre to centre), also called nucleosome repeat length or NRL (*i*.*e*., the length of DNA wrapped around the histone octamer plus linker DNA), depending on the considered species and culture condition. In *B. cinerea*, average NRL is estimated at 169 bp (Figure 1A). In *F. graminearum*, this distance reaches 172 bp (Figure 1B). In Lmb and Lml (Figures 1C and 1D), average NRL is 166 bp and 161 bp, respectively. Considering these values and the canonical length of nucleosomal DNA (147 bp), linker DNA length can be estimated to stand, in average, between 14 to 19 bp for respectively Lml and Lmb, 22 bp for *B. cinerea*, and 25 bp for *F. graminearum*. Nucleosome phasing genome-wide seems to be particularly tight in *F. graminearum*, with very little deviation in the measured phases (Figure 1B and Supplementary Table 2). In contrast, higher deviations are observed for *B. cinerea* (Figure 1A and Supplementary Table 2).

**Figure 1:**
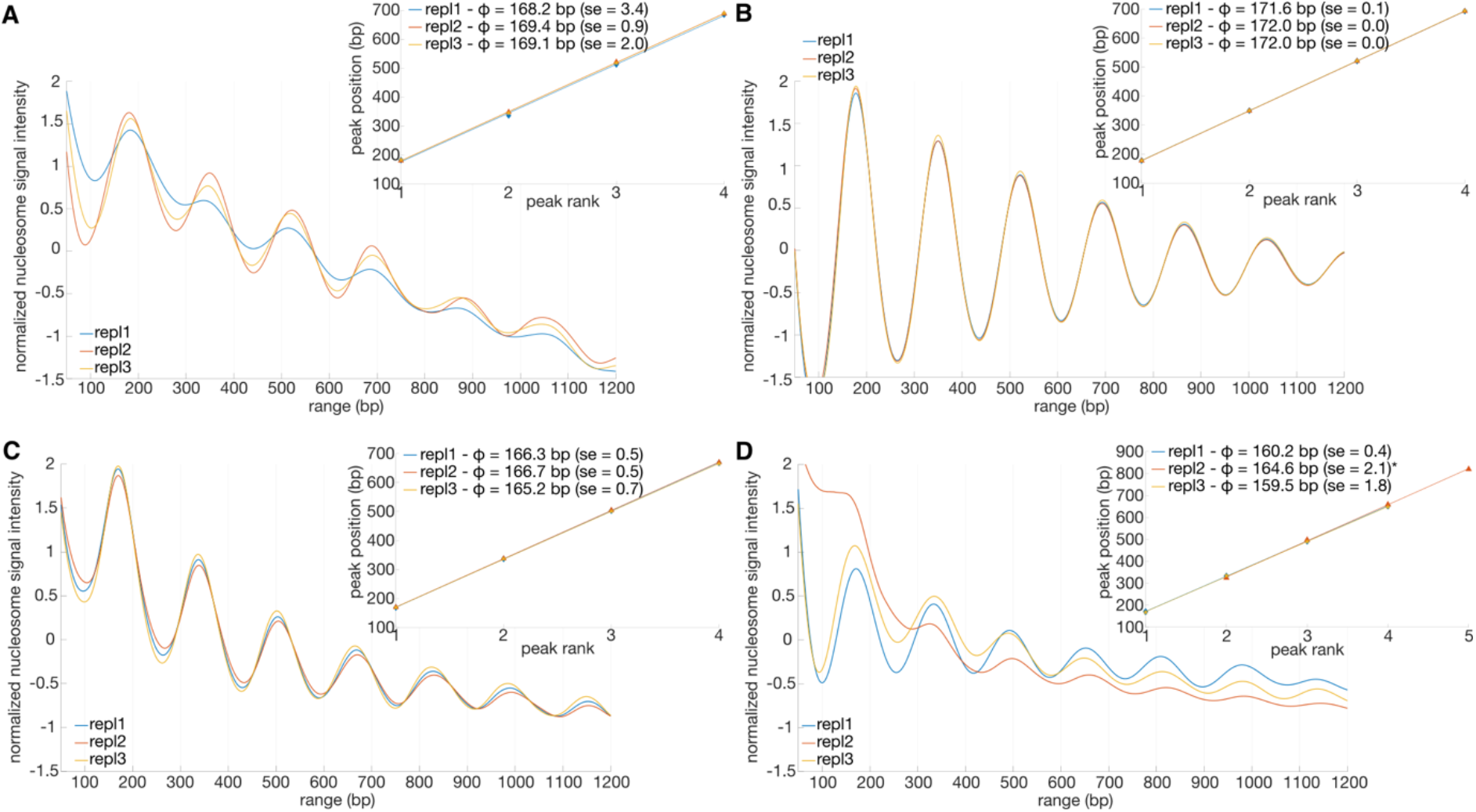
Nucleosome phasing in the four fungi studied. Main graphs display scaled (z-score) phase frequencies (y-axis) as a function of position (in base pair; x-axis). Graphs in inserts show peak positions (in base pairs; y-axis) as a function of peak order (x-axis). For each replicate, phases, standard errors (se), R^2^ (coefficient of determination), and *p*-values (*F*-test) are determined after linear regression fitting to the first four successive peak positions (see Supplementary Table 2). Repl = replicate. A. *Botrytis cinerea*; B. *Fusarium graminearum*; here, phase value in replicate #2 was measured for four successive peaks excluding peak #1 for which an apex was not clearly visible at the beginning of the profile; C. *Leptosphaeria maculans* ‘brassicae’; D. *Leptosphaeria maculans* ‘lepidii’.

Nucleosome spacing influences the formation of the higher order chromatin fibre, often referred to as the 30-nm chromatin fibre (Szerlong and Hansen, 2011). Several structural models of the chromatin fibre have been proposed, all underlining nucleosome-nucleosome interactions including the length of linker DNA fragments as major driving factors (Zhu and Li, 2016). Notably, the chromatin fibre was found to be narrower (21-nm in diameter) for a short NRL of 167 bp (Robinson *et al*., 2006). Similarly, increasing NRLs lead to increasingly wider fibres, reaching a highly compact 30-nm solenoid structure for an NRL of 197 bp. Typically, NRLs are ∼175-200 bp in plants, *Caenorhabditis elegans*, and humans (Valouev *et al*., 2008, 2011; Locke *et al*., 2013; Zhang, Zhang and Jiang, 2015), and ∼165 bp and 154 bp in the yeasts *S. cerevisiae* and *Schizosaccharomyces pombe*, respectively (Yuan *et al*., 2005; Lantermann *et al*., 2010). In the present study, we found NRL values remarkably constant between biological replicates, a phenomenon sometimes referred to as clamping that involves ATP-dependent chromatin remodelers in purified experimental systems (Lieleg *et al*., 2015). The species *B. cinerea* and *F. graminearum* show similar NRLs in the middle range (169 bp to 172 bp), possibly indicating intermediate levels of compaction of the higher order chromatin fibre. These values are similar to those obtained for the Pezizomycotina *A. fumigatus, i*.*e*., linker length ranging from 21 to 27 bp, using an MNase treatment similar to the one used in the present study (Nishida *et al*., 2009). Lml and Lmb distinguish themselves with shorter NRLs of only 161-166 bp, suggesting a narrower chromatin fibre structure. In the yeast *S. cerevisiae*, linker length was shown to be the result of the competition for binding between the chromatin remodeling factors ISW1a (Imitation SWItch) and CHD1 (Chromodomain Helicase DNA-binding), the latter mediating shorter length after the eviction of histone H1 (Ocampo *et al*., 2016). Considering that sequence polymorphisms in CHD1 has been previously associated with variations of linker length (Hughes and Rando, 2015), a seducing possibility is that such variation at the protein level may account for a portion for the inter-species differences observed here. Regarding the very closely related species Lmb and Lml, Lmb presents longer NRLs than Lml. We hypothesized this peculiarity may be explained by large AT-rich regions displayed by the Lmb genome, not encountered in the Lml genome (Rouxel *et al*., 2011; Grandaubert *et al*., 2014). Indeed, DNA sequence is a major determinant of nucleosome landscapes (Struhl and Segal, 2013), in particular AT stretches that confer more rigidity to the chromatin fiber.

### Nucleosome distribution profiles

Read density was plotted genome-wide in one kb-long sliding non-overlapping windows along chromosomes for all four fungi (Figures 2 to 4, and Supplementary Figures 2 to 6). The density profiles obtained for *B. cinerea* show remarkable regularity of nucleosome density genome-wide (Figure 2A and Supplementary Figure 2). Nevertheless, we could observe that almost all occasional thin peaks of density were correlated with the positions of BOTY retro-transposons (Porquier *et al*., 2016). Out of 48 complete copies of BOTY in the genome, 31 show a peak of nucleosome density. Notably, they correspond to the BOTY elements with an equilibrated percentage of GC (43-45%) while the 17 copies that do not show such a peak are those with a lower percentage of GC (14-24%) probably because they have undergone Repeat-Induced Point mutation, or RIP (Porquier *et al*., 2016). Peaks of density were rarely observed for TE other than BOTY (Supplementary Figure 3). We also investigated nucleosome spacing in regions occupied by BOTY and non-BOTY TE. Phasograms were plotted as described above restricting our analysis to BOTY or other TE (Figures 2B and 2C, respectively). Much larger phases can be observed in other TE regions (178.4-187.5 bp) when compared to BOTY regions (171.3-172.5 bp) or genome-wide (168.2-169.4 bp, Figure 1A), indicating larger nucleosome spacing in TE-occupied regions. BOTY-containing regions, which positions correlate with discrete peaks of nucleosome density, exhibit slightly larger phases than genome-wide. Thus, the observed peaks of read density may be the result of increased nucleosome occupancy, *i*.*e*., a measure of the stability of a nucleosome at a given position in a multiple cell sample, rather than a denser deposition of nucleosomes. BOTY is one of the largest TE identified in the B05.10 strain (6.4-6.6 kb), and that’s definitively the TE with the largest genome coverage *i*.*e*., 0.96% (Porquier *et al*., 2016). Notably, the majority of *B. cinerea* small interfering RNA (siRNA) predicted to silence host plant genes are derived from the copies of BOTY and related elements that show an equilibrated percentage in GC (Weiberg *et al*., 2013; Porquier *et al*., 2021). As the production of these siRNA effectors is activated during the early phase of plant infection, we could speculate that the high nucleosome occupancy on the *loci* of production (*i*.*e*. un-RIPped BOTY TEs) is a mechanism to restrict their production during saprophytic growth. The observation of two distinct chromatin states characterizing TEs in *Verticillium dahliae* supports this proposition (Cook *et al*., 2020). This hypothesis remains to be tested by investigating nucleosome occupancy during *in planta* development.

**Figure 2:**
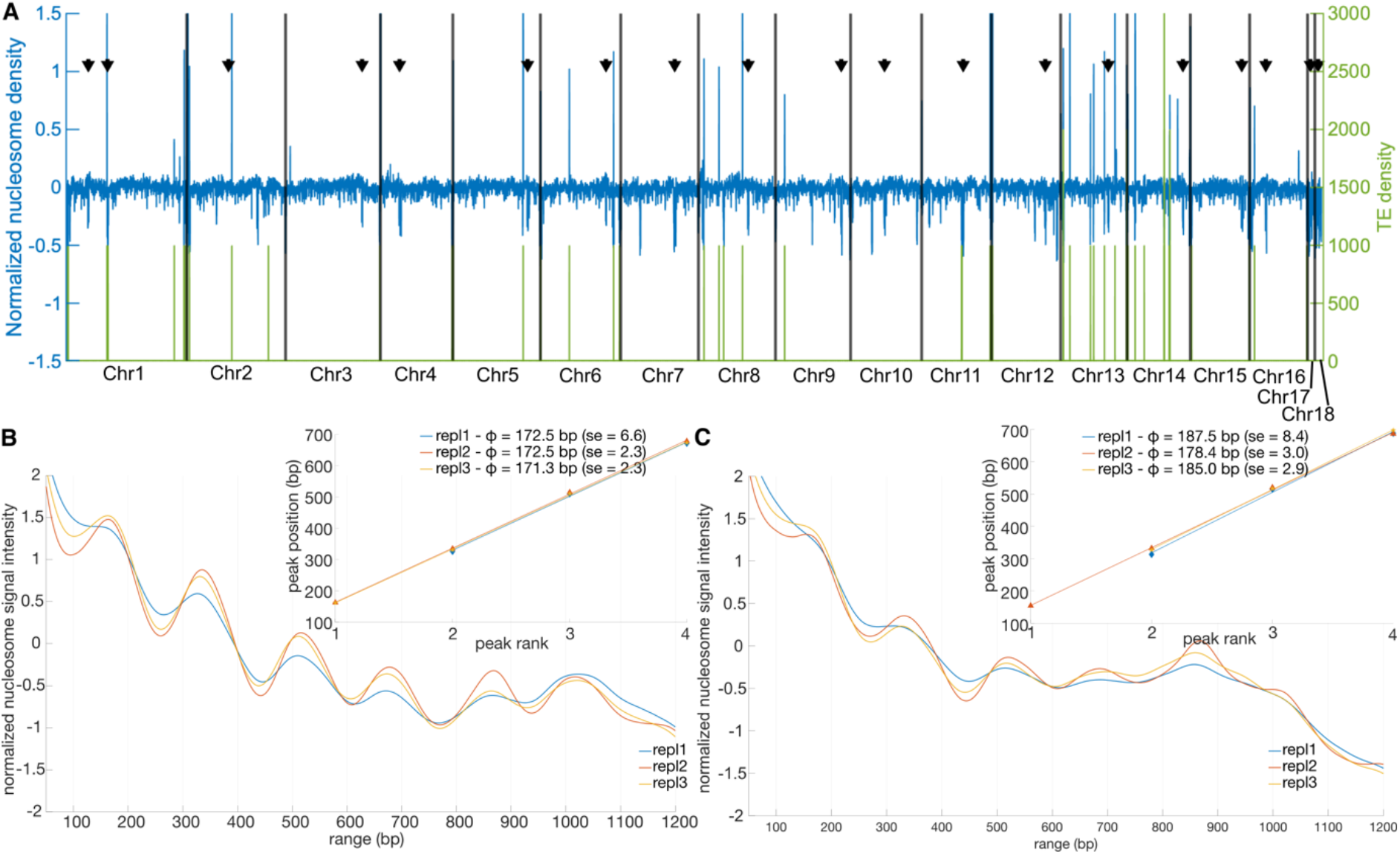
Nucleosome density profiles in *Botrytis cinerea*. A. Coverage density profiles were computed for non-overlapping 1 kb-long bins along the chromosomes of B. cinerea. In green are plotted BOTY transposable elements (TE) density profiles. In blue are plotted the z-scored average nucleosome density profile (see Supplementary Figure 2 for individual replicate plots). Black arrows indicate putative positions of centromeres (Van Kan *et al*., 2017); B and C. Nucleosome phasing in BOTY TE (B) or TE other than BOTY TE (C). Main graphs display phase frequencies (y-axis) as a function of position (in base pair; x-axis). Graphs in inserts show peak positions (in base pairs; y-axis) as a function of peak order (x-axis); Phases +/- standard errors (se), R2 (coefficient of determination), and p-values (F-test) are determined after linear regression fitting to the first four successive peak positions (see Supplementary Table 3). Repl= replicate.

In *F. graminearum*, regions equally packed with nucleosomes are interspaced with areas with lower density (Figure 3A and Supplementary Figure 4). Strikingly, this profile mirrors the previously described SNP density profiles in *F. graminearum* (Laurent *et al*., 2017). We investigated whether or not this profile was the result of increased spacing between nucleosomes in regions found denser in SNPs. Phasograms were plotted restricting our analysis to the SNP-enriched polymorphic islands or the rest of the genome, as defined by Laurent et al. (Laurent *et al*., 2017). Wave signals similar to the ones observed genome-wide were obtained, with phases of 172.3-172.4 bp in polymorphic islands (Figure 3B, Supplementary Table 3) and 171.6-171.9 bp outside these regions (Figure 3C, Supplementary Table 3). These results indicate that nucleosomes appear well-arrayed genome-wide, with very similar phases in polymorphic islands *vs*. non-polymorphic islands. Thus, the observed drops in read density profile cannot be explained by a depletion in nucleosomes but may rather be the result of reduced nucleosome occupancy. This observation suggests increased frequencies of transient nucleosome positioning events in *F. graminearum* fast evolving polymorphic islands (Laurent *et al*., 2017) and thus more relaxed chromatin structures. Here, nucleosome dynamics may enable fast evolution of particular genome segments while regions defined by higher occupancies may secure sequence conservation.

**Figure 3:**
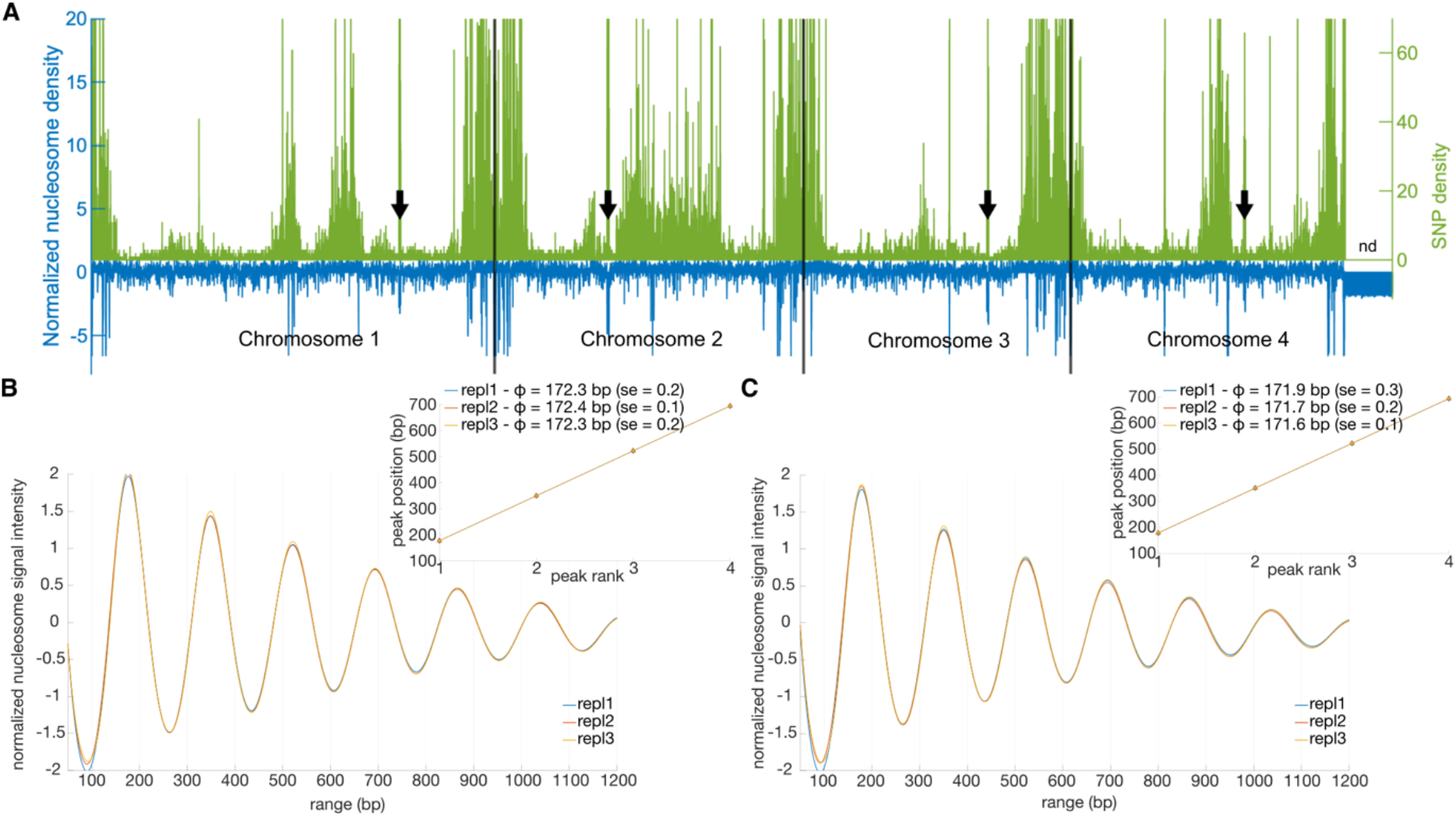
Nucleosome density profiles in *Fusarium graminearum*. A. Coverage density profiles were computed for non-overlapping 1 kb-long bins along the four chromosomes of *F. graminearum*. In green are plotted SNP density profiles as previously described (Laurent *et al*., 2017). “nd” indicates the highly variable 3’ end of chromosome 4 for which SNP were not called. In blue are plotted the z-scored average nucleosome density profile (see Supplementary Figure 2 for individual replicate plots). Black arrows indicate centromeres (King *et al*., 2015); B and C. Nucleosome phasing in polymorphic islands (B) or outside polymorphic islands (C) as previously defined (Laurent *et al*., 2017). Main graphs display phase frequencies (y-axis) as a function of position (in base pair; x-axis). Graphs in inserts show peak positions (in base pairs; y-axis) as a function of peak order (x-axis); Phases +/- standard errors (se), R^2^ (coefficient of determination), and *p*-values (*F*-test) are determined after linear regression fitting to the first four successive peak positions (see Supplementary Table 3). Repl = replicate.

In Lmb, numerous “islands” of nucleosome-dense regions can be observed at various locations of the genome, including the dispensable chromosome (Figure 4A and Supplementary Figure 5). Aside from couple of contigs displaying higher nucleosome density, such characteristics were not observed for the closely related species Lml (Supplementary Figure 6). The locations of these nucleosome-dense islands in Lmb parallel those of AT-rich regions of the genome (Figure 4A and Supplementary Figure 5), features not visible in the genome of Lml (Supplementary Figure 6), suggesting that AT-rich regions are particularly dense in nucleosomes. Considering the remarkable compartmentalized organization of the genome of Lmb (Rouxel *et al*., 2011), absent from Lml, differences of nucleosome phasing and occupancy in TE- and AT-rich *vs*. GC-equilibrated and gene-rich regions were investigated. A region was considered AT-rich if it contained less than 40 % of GC. As described earlier, AT-rich regions represent one-third of the Lmb genome divided in 419 regions of 1 to 320 kb in length. Examination of unprocessed mapping outputs reveals that the number of fragments (read pairs) mapped in AT-rich and GC-equilibrated regions were very similar, with 23.8 million and 24.8 million fragments, respectively, which is far from the 1/3 *vs*. 2/3 ratio expected. In terms of coverage depth, mean coverage is 207 *vs*. 135 fragments for AT-rich and GC-equilibrated regions, respectively, which could suggest higher nucleosome occupancy in the former. We explored this hypothesis and compared phasograms for AT-rich *vs*. GC-equilibrated regions (Figure 4B and 4C, Supplementary Table 3). Average NRLs were found larger in AT-rich than GC-equilibrated compartments, measured at 169.2 bp and 164.2 bp respectively, suggesting lower nucleosome frequencies in AT-rich regions than in GC-equilibrated regions. Nonetheless, coverage density is higher in AT-rich regions (Figure 4A and Supplementary Figure 5), consistently with our hypothesis of higher nucleosome occupancy in these regions and thus less accessible genome compartment, in heterochromatic state. This is in accordance with the recent genome-wide mapping of histone modifications performed by Soyer et al. (2020) on Lmb and Lml in which the Histone H3 Lysine9 tri-methylation heterochromatin mark was found associated with TE- and AT-rich regions of Lmb including on the dispensable chromosome (Soyer *et al*., 2020). Finally, signal intensity in phasograms appeared more stable on the long nucleotide range in GC-equilibrated than in AT-rich regions, an observation in line with the well-known destabilizing effect of AT stretches on nucleosome positioning leading to fuzzier signals (Kaplan *et al*., 2009; Ponts *et al*., 2010; Tillo *et al*., 2010; Bunnik *et al*., 2014; Russell *et al*., 2014; Jin, Finnegan and Song, 2018). All together, these data support a heterochromatic state of Lmb AT-rich regions mediated by strong nucleosome occupancy during axenic growth. Since the AT-rich regions host many fungal effector genes expressed during primary infection of oilseed rape leaves, we may assume that these regions are decondensed during infection, allowing the action of specific transcription factors. We tried to perform MNase-seq experiments at an early stage of oilseed rape infection by Lmb but the number of fungal reads was too low to be able to reliably analyze fungal nucleosome positioning. To go further, techniques to enrich in fungal material prior to MNase treatment should be considered.

**Figure 4:**
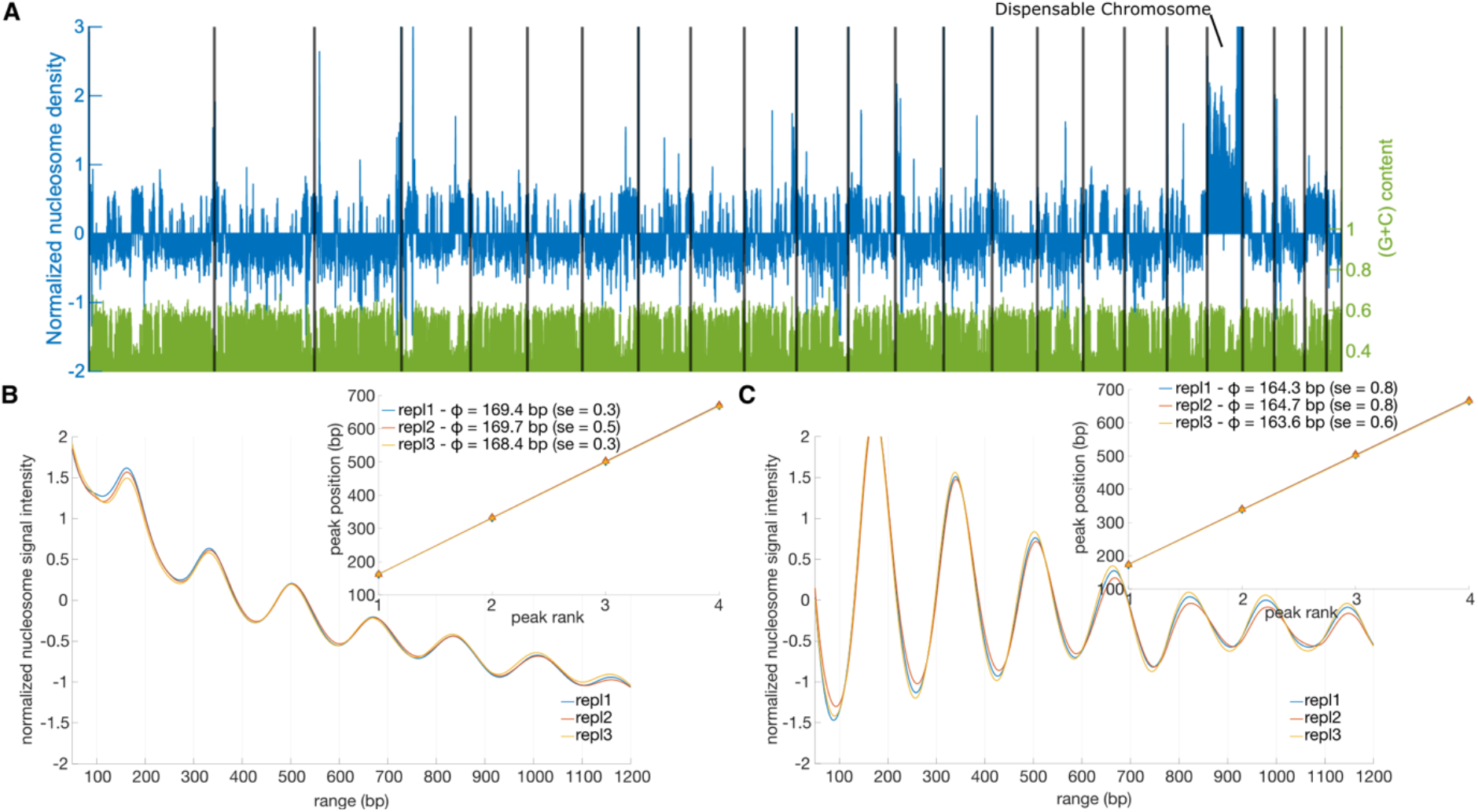
Nucleosome density profiles in *Leptosphaeria maculans* ‘brassicae’. A. Coverage density profiles were computed for non-overlapping 1 kb-long bins along all supercontigs, separated by black lines. (G+C)/(A+T+G+C) content is plotted in green. In blue are plotted the z-scored average nucleosome density profile (see Supplementary Figure 3 for individual replicate plots). B and C. Nucleosome phasing in TE and AT-rich regions previously described in (Rouxel *et al*., 2011) (B) and GC-equilibrated regions (C). Main graphs display phase frequencies (y-axis) as a function of position (in base pair; x-axis). Graphs in inserts show peak positions (in base pairs; y-axis) as a function of peak order (x-axis); for each replicate, phases, standard errors (se), R^2^ (coefficient of determination), and *p*-values (*F*-test) are determined after linear regression fitting to the first four successive peak positions (see Supplementary Table 3). Repl = replicate.

### Sequence composition and nucleosome positioning

Literature data report that distribution of bases in nucleosome core DNA is non-random and exhibits a ∼10 bp AA/TT/AT/TA offset with GG/CC/GC/CG dinucleotide frequency (Satchwell, Drew and Travers, 1986; Segal *et al*., 2006; Segal and Widom, 2009b). Here, we investigated di-nucleotide frequencies – *i*.*e*., the incidence of a given neighbouring pair of nucleotides in a sequence – in nucleosomal DNA segments in all four fungi. Averaged di-nucleotide contents centred around all fragments (read pairs) were plotted (Figure 5) and periodicities investigated by autocorrelation analyses (Supplementary Figure 7). Autocorrelation plots reveal the previously described ∼10 bp-periodicities (Laurent *et al*., 2017)(Satchwell, Drew and Travers, 1986; Segal *et al*., 2006; Segal and Widom, 2009b) for all studied fungi while showing differences in instant autocorrelation coefficient profiles (Supplementary Table 4 and Supplementary Figure 7). Signal is indeed very regular in *F. graminearum* and, to a lesser extent, Lmb, whereas Lml and *B. cinerea* show more irregular autocorrelation profiles. These results are consistent with our previous observation that nucleosomes are tightly phased in *F. graminearum* whereas somehow fuzzier (higher deviation) in *B. cinerea* and Lml (Figure 1A and 1D).

**Figure 5:**
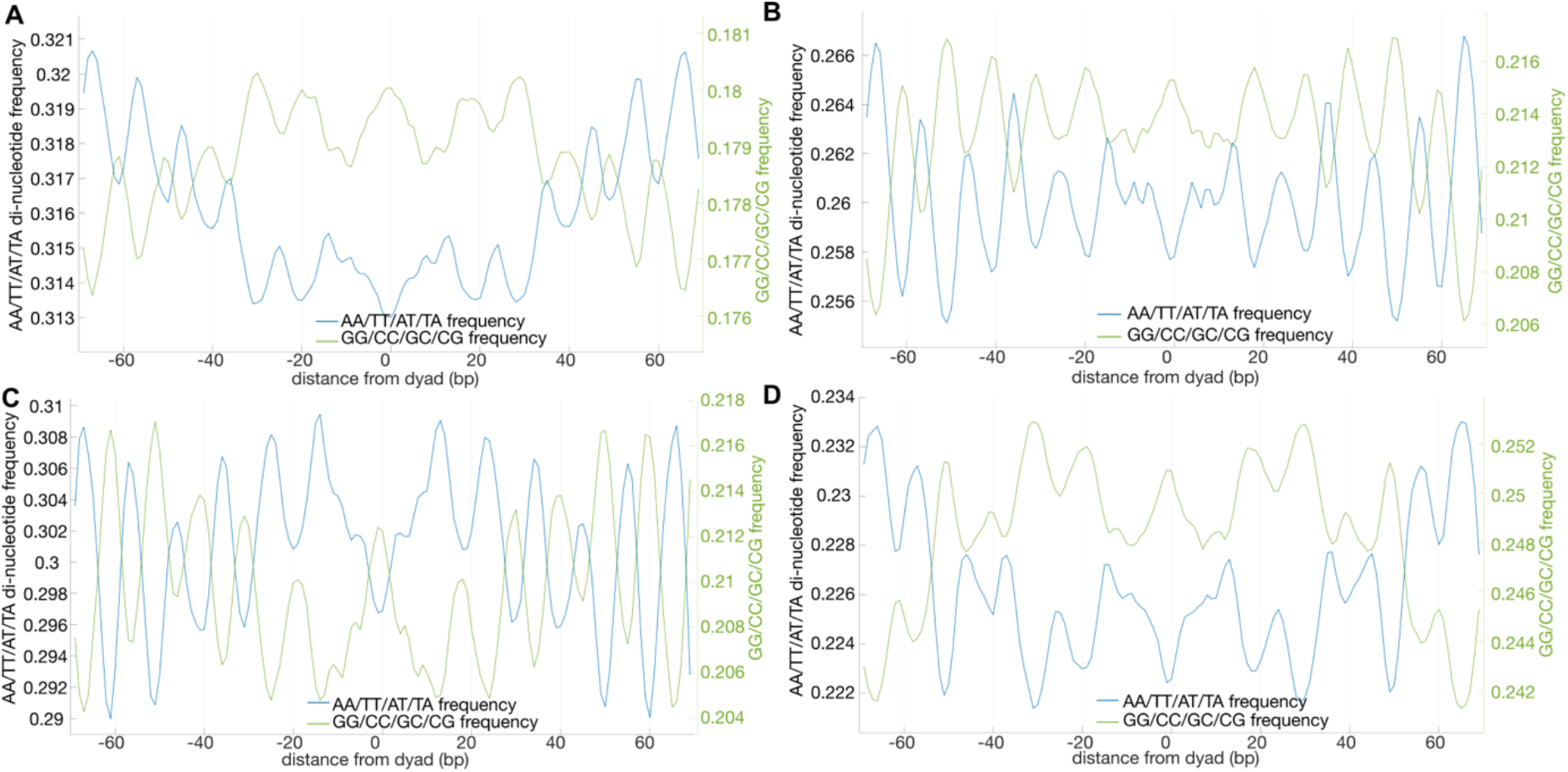
Repeated di-nucleotide patterns in nucleosomal DNA. Di-nucleotides frequency plots (average of three biological replicates) for *Botrytis cinerea* (A), *Fusarium graminearum* (B), *Leptosphaeria maculans* ‘brassicae’ (C), and *Leptosphaeria maculans* ‘lepidii’ (D).

Di-nucleotide frequency graphs display A+T dinucleotides frequency waves oscillating out of phase with G+C ones. For all studied fungi, GC dinucleotides are centred on nucleosome dyads (Figure 5). Considering that there are 16 possible combinations of di-nucleotides, equilibrated distribution of di-nucleotide contents should contain 25% of AT/TA/AA/TT and 25% of GG/CC/CG/GC. Our observations reveal a skewed distribution in favour of AT dimers marked for *B. cinerea* and Lmb (Figure 5A and 5C. The presence of AT-rich regions in Lmb and, to a lesser extent, in *B. cinerea* genomes may explain such a result (Rouxel *et al*., 2011; Porquier *et al*., 2016). Di-nucleotide frequencies of Lmb AT-rich and GC-equilibrated regions were thus inspected (Supplementary Table 5 and Supplementary Figure 8). As one would expect, A+T frequencies are particularly high in AT-rich regions, while maintaining alternance with G+C di-nucleotides and ∼10 bp periodicity.Nucleosome positioning is believed to be particularly hard-wired to DNA sequence, and especially the largely documented anti-nucleosome effect of Poly(dA:dT) tracts (Segal and Widom, 2009a; Struhl and Segal, 2013). Here, while nucleosome phase was indeed found 5 bp longer in Lmb AT-rich regions than in GC-equilibrated regions, occupancy was nonetheless higher in the former leading to the formation of the previously suggested heterochromatic state of these regions, which has consequences on gene expression and recombination (Rouxel *et al*., 2011; Soyer *et al*., 2014, 2020; Gay *et al*., 2020). These observations suggest the mobilisation of trans-acting chromatin remodeling factors to maintain heterochromatin structures on such disfavouring sequences. Importantly, we found that GC periodicity at nucleosome dyads is preserved even within AT-rich regions, suggesting such pattern in an AT-rich environment is sufficient to permit efficient wrapping of DNA around nucleosomes and strong occupancy. This longer NRLs in addition with high nucleosome density in AT-rich regions should have an impact on gene expression in these regions enriched in effector gene (Rouxel *et al*., 2011; Soyer *et al*., 2020).

### Nucleosome landscapes of fungal gene units

Nucleosome occupancy profiles of gene units were investigated in all fungi (Figure 6). As previously reported in other eukaryotes, translation start sites (the ATG codon) are preceded by a nucleosome-depleted region (NDR) and immediately followed by a well-positioned +1 nucleosome (Figure 6A). Variations of the exact position of these features relative to the ATG start codon, as well as variations in the intensity of the NDR valley, are nonetheless observed between fungal species. For example, the NDR and the centre of the +1 nucleosome (or +1Nucl) are found at -154 bp and +14 bp, respectively, in *F. graminearum* whereas they are located at -99 bp and + 26 bp, respectively, in *B. cinerea*. The fungus *B. cinerea* shows the deepest NDR valley among all observed profiles. In Lmb and Lml, NDRs are found at - 129 and -144 bp from ATG, respectively, and +1Nucl at +19 bp and +55 bp. Finally, nucleosome profiles upstream of NDRs appear fuzzy for all fungi but Lmb, with varying degrees of fuzziness. This fuzziness is no longer visible when nucleosome profiles are centred on TSS for *F. graminearum* (N_TSS_ = 6,212 genes) and *B. cinerea* (N_TSS_ = 11,701 genes) (Figure 6C). The NDR is more defined, and located immediately upstream of the TSS, with a minimum detected at -58 bp and -20 bp upstream of the TSS of *F. graminearum* and *B. cinerea*, respectively. These values are consistent with the binding of the RNA polymerase II ∼50 bp upstream of TSS, observed in active promoters of mammalian and Drosophila cells (Core, Waterfall and Lis, 2008; Min *et al*., 2011; Kwak *et al*., 2013).

**Figure 6:**
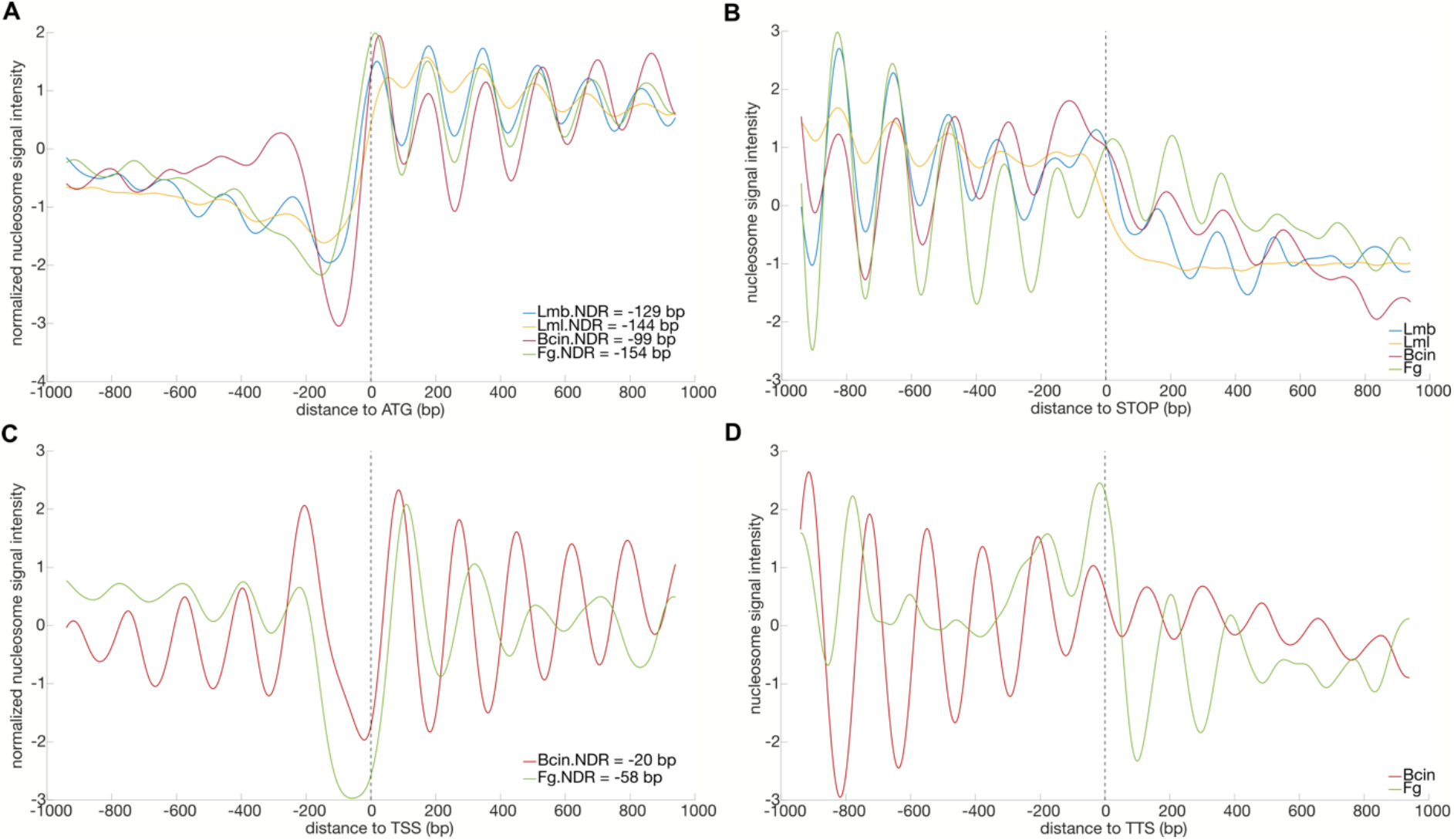
Nucleosome organization of fungal gene units. A-D. Scaled (z-scored) averages (three biological replicates for each fungus/condition) of nucleosome signals as a function of position (in base pairs) relative to the start codon ATG (A), the stop codon (B), TSS (C), TTS (D). Fg = *Fusarium graminearum*; Lmb = *Leptosphaeria maculans* ‘brassicae’; Lml = *Leptosphaeria maculans* ‘lepidii; Bcin = *Botrytis cinerea*; NDR = nucleosome-depleted region.

Considering nucleosome environments at stop codons (Figure 6B), strongly arrayed nucleosomes are particularly found upstream the stop codon, fewer signal variations being observed downstream. Here, the stop codon is a clear boundary for nucleosome arraying and occupancy in all fungi and all conditions investigated. A nucleosome seems remarkably well positioned exactly on stop codons in *F. graminearum* in particular. The analysis was repeated on TTS in *F. graminearum* (N_TTS_ = 5,292 genes) and *B. cinerea* (N_TTS_ = 11,701 genes) (Figure 6D). In *B. cinerea*, signal appeared strong and well-arrayed, decaying downstream of the TTS. In *F. graminearum*, strong positioning of nucleosomes -2 and -1 (−178 bp and -16 bp relative to TTS, respectively) followed by a deep 3’ end NDR (+100 bp downstream of TTS) can be observed. Nucleosome positioning on TSS and TTS could not be analyzed for *Lmb* and *Lml* since TSS and TTS annotations are not supported by experimental data for these species’ genomes as it is the case for *F. graminearum* (King, Urban and Hammond-Kosack, 2017; Basenko *et al*., 2018), or by collaborative annotation as for *B. cinerea* (Pedro *et al*., 2019).

For comparison purposes, nucleosome profiling was repeated restricting our analysis to genes identified as BUSCO lineage-specific single-copy evolutionary conserved orthologs in Fungi (Simão *et al*., 2015; Waterhouse *et al*., 2018). Overall broad patterns remain similar to those obtained while investigating whole genomes, with notably somehow more regular oscillations patterns (Figure 7). In *F. graminearum*, whilst the distance NDR-to-TSS (− 60 bp) remains very similar to the one measured earlier genome-wide (Figure 7C), distance NDR-to-ATG increases by 29 bp (Figure 7A), whereas in *B. cinerea* the distance NDR-to-TSS reduces to 0 bp and nearly no increase in the distance NDR-to-ATG is observed (+ 3 bp). Similarly, the distance NDR-to-ATG is only 5 bp longer than genome-wide in Lmb whereas it increases by 23 bp in Lml. Towards the 3’ end of the gene unit, nucleosome signals around stop codons are similar to those obtained genome-wide for all fungi (Figure 7B). However, the deep NDR found downstream of TTS of *F. graminearum* genome-wide can no longer be observed at “Fungi” TTS *loci* (Figure 7D). Similar to genome-wide profile, strongly positioned -1Nucl and -2Nucl are still visible at -5 bp and -209 bp, respectively, the latter being more intense and defined.

**Figure 7:**
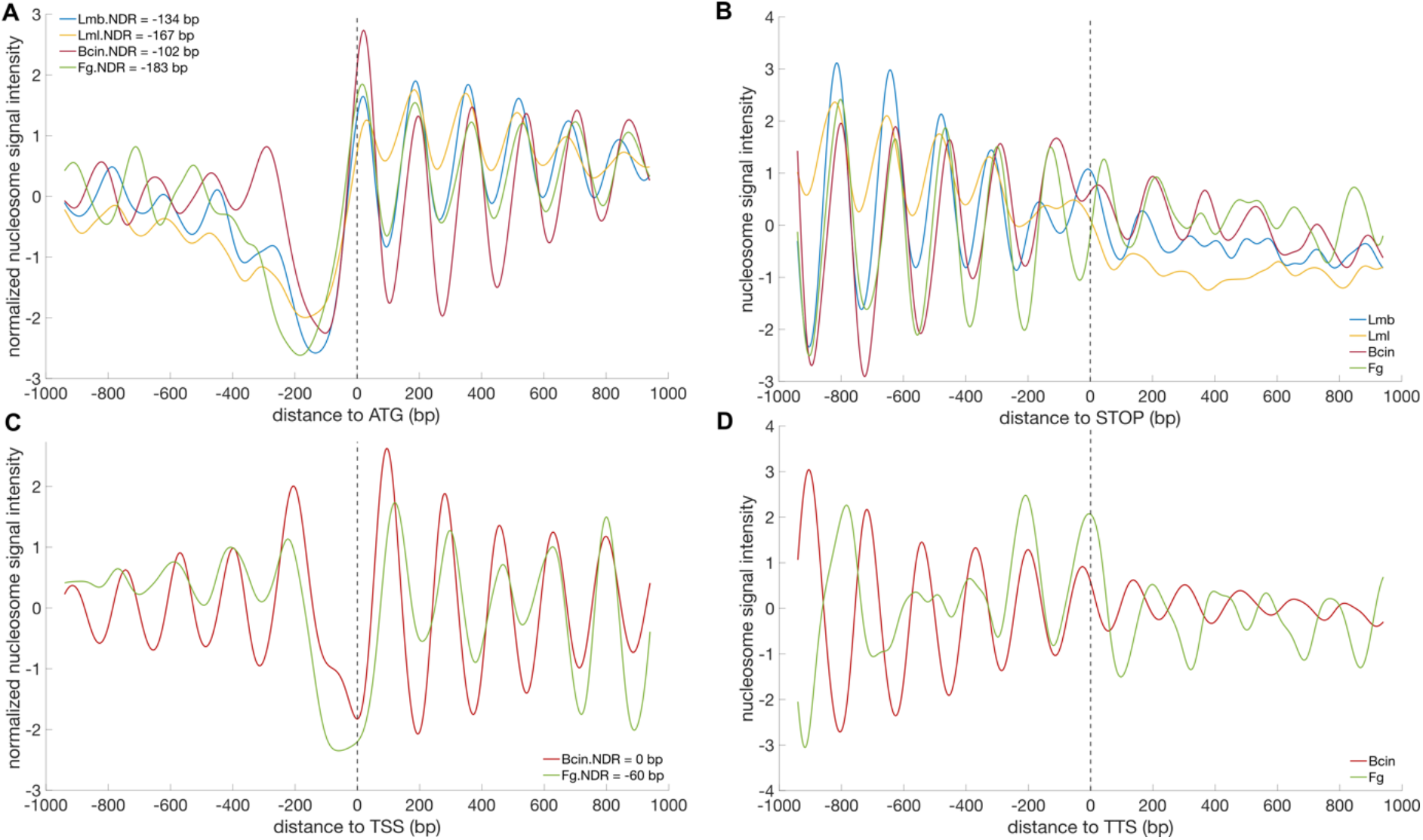
Nucleosome organization of near-universal single copy orthologous gene units in Fungi (BUSCO3). A-D. Scaled (z-scored) averages (three biological replicates for each fungus/condition) of nucleosome signals as a function of position (in base pairs) relative to the start codon ATG (A), the stop codon (B), TSS (C), and TTS (D). Fg = *Fusarium graminearum*; Lmb = *Leptosphaeria maculans* ‘brassicae’; Lml = *Leptosphaeria maculans* ‘lepidii; Bcin = *Botrytis cinerea*; NDR = nucleosome-depleted region.

The general profile of a fungal gene unit shares similarities with those previously described in various eukaryotes: the ATG codon is decorated by a well-positioned +1 nucleosome and preceded by an NDR. This “+1 nucleosome” is an extremely well-conserved feature among eukaryotes spread across the tree of life, and nucleotide sequence only is not sufficient to explain such consistency. Such stability requires the intervention of ATP-dependent chromatin remodelers, belonging to one of the families CHD, INO80, ISWI, or SWI/SNF (Reyes, Marcum and He, 2021). Nucleosome landscapes can thus be viewed as the final result of active positioning forces (the action of chromatin remodelers) combined with destabilizing nucleotide content, including poly(A+T) tracts (see above). Recently, this scenario was proposed to be species-specific (Barnes and Korber, 2021; Oberbeckmann, Krietenstein, *et al*., 2021), supporting that several combinatorial nucleosome arraying rules can form during the course of evolution. Indeed, when we restricted our analysis to conserved single-copy orthologous fungal genes, the overall profiles and the intensities of “+1 nucleosome” and NDRs were more homogenous between fungi. Similarly, while an NDR can be observed downstream of *F. graminearum* TTS, it is no longer visible when the analysis is restricted to conserved fungal genes, indicating again an evolutionary component. Promoters are typically found in NDRs upstream of +1 nucleosomes (Yuan *et al*., 2005; Jiang and Pugh, 2009). NDR sizes largely depend on the action of the SWI/SNF ATP-dependent remodeler RSC (Remodeling the Structure of Chromatin) complex (Krietenstein *et al*., 2016; Yague-Sanz *et al*., 2017; Wagner *et al*., 2020) that seem to facilitate initiation of transcription by preventing the filling of NDRs with nucleosomes (Ocampo *et al*., 2019).

### Nucleosome landscapes of gene units according to gene expression

Same analyses were repeated for genes categorised according to their expression levels (expressed in TPM counts, see Materials and Methods). The general variations in nucleosome profiles around translation start sites are similar for all expression categories in all considered fungi and culture conditions: ATG codons are immediately followed by a well-positioned +1 nucleosome and preceded by a dip in nucleosome density (Figure 8). Remarkable variations are nonetheless observed with regard to positions of +1 nucleosomes and NDRs, as well as the amplitude of the nucleosome signal difference between them (here defined as Δnucl = |signal_+1nucl_ -signal_NDR_|), depending on gene expression. ATG-centred nucleosome profiles for genes not expressed in our conditions (TPM = 0) show remarkably reduced Δnucl when compared to those measured for genes more expressed, and a distance to the ATG reduced (Figure 8 and Supplementary Table 6). Conversely, highly expressed genes (TPM50) display the deepest NDRs located at the furthest upstream the ATG. Similar trends are observed when profiles are centred on *B. cinerea* and *F. graminearum* TSS (Figures 8E and 8F, respectively). Moreover, NDRs were usually found further from the ATG site than those in genes expressed at lower levels or not expressed. Finally, the nucleosome wave signal decay phenomenon was observed at distances from the ATG codons shorter in poorly expressed genes than in highly expressed genes, although the +1 nucleosome remained fairly well conserved.

**Figure 8:**
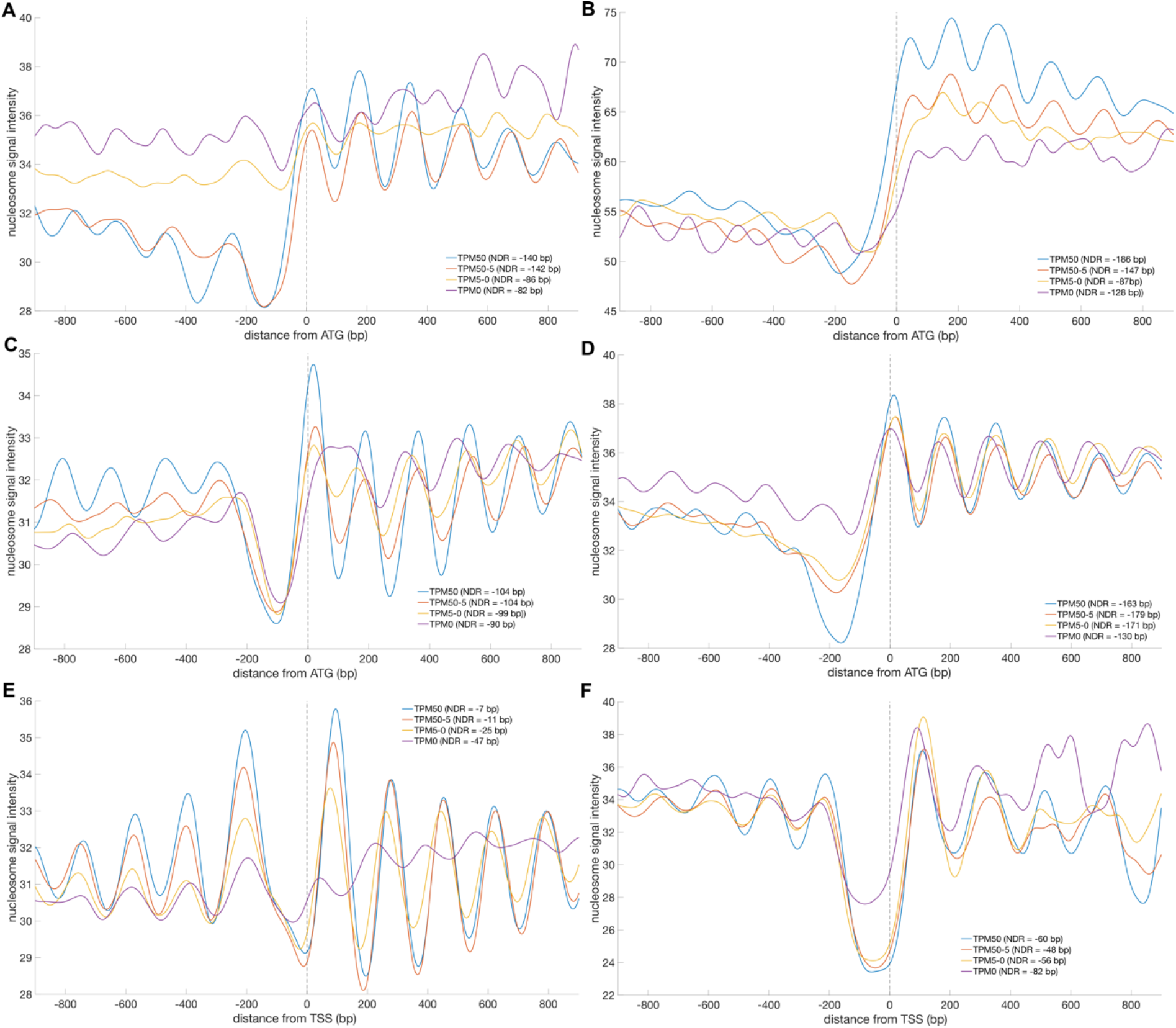
Nucleosome organization at start codons/TSS *vs*. gene expression. Average (three biological replicates for each fungus/condition) nucleosome signal as a function of position (in base pairs) relative to the start codon ATG (and TSS for *Fusarium graminearum* and *Botrytis cinerea*). TPM = Transcripts Per Million. (A), In *Leptosphaeria maculans* ‘brassicae’, ATG-centred; (B), In *Leptosphaeria maculans* ‘lepidii’, ATG-centred; (C), In *B. cinerea*, ATG-centred; (D), *F. graminearum*, ATG-centred; (E). In *B. cinerea*, TSS-centred; (F). *F. graminearum*, TSS-centred.

Nucleosome depletion can be the side-effect of active transcription with the binding of pre-initiation complex resulting in nucleosome eviction as previously shown in yeast (Venters and Pugh, 2009). Indeed, we evidenced valleys of nucleosome signals upstream of ATG codons found deeper in highly expressed genes. On the contrary, genes not expressed showed little or no NDR, depending on the considered species or culture condition. In our conditions, the amplitude between the NDR and the +1 nucleosome seemed to be an informative measure of gene expression level: the higher this value is, the more genes are expressed. This feature was less strict when TSS were considered, raising the question of different mechanisms of transcription regulation depending on gene unit structures. A possible scenario is that other factors, such as the general regulator Reb1 (RNA polymerase Enhancing Binding protein), that modulate the action of remodeling factors (Ghassabi Kondalaji and Bowman, 2021) may contribute to the profiles observed here. Strikingly, a recent study evidenced the role of such factors as barriers to fine-tune the action of remodelers, with the consequence of modulating nucleosome spacing and phasing distances (Oberbeckmann, Niebauer, *et al*., 2021). Oberbeckmann and Colleagues proposed a model consisting in promoter NDRs (maintained by the RSC complex, see above) insulated upstream by the barrier factor Reb1 and downstream by the +1 nucleosome, the spacing between the two landmarks being controlled by the remodelers INO80 or ISW2, maintaining longer *vs*. shorter distances respectively (Oberbeckmann, Niebauer, *et al*., 2021). In this model and consistently with our observations, gene bodies are characterized by high nucleosome density with shorter NRLs. Here, the marked oscillations of the signal obtained for highly expressed genes suggests that the presence of well-arrayed nucleosomes around ATG combined with high amplitudes and longer distances of nucleosome signal between the NDR valley and the +1 nucleosome peak could be a hallmark of robust active gene expression. On the contrary, an absent or virtually absent NDR upstream of ATG may mark genes with variable expression levels, often exhibiting a TATA-box in their promoters (Tirosh and Barkai, 2008). Such promoters displayed enhanced sensitivity to mutations in yeast, an observation arguing for a link between chromatin structure and the evolution of gene expression (Hornung, Oren and Barkai, 2012). A further confounding element is the observation that introducing Reb1 binding sites in such promoters reduced sensitivity to mutation (Hornung, Oren and Barkai, 2012), which was interpreted in 2012 (when the work was published) as blocking nucleosome formation and introducing an NDR. In the light of the model recently proposed by Oberbeckmann *et al*., (2021; see above), this analysis may now be re-visited as providing the necessary binding factor that constraints NDR formation and maintenance by RSC and INO80/ISW2.

At the end of gene units, wave signal decay was also observed downstream of stop codons in all studied fungi (Supplementary Figure 9). In Lmb and Lml, strong nucleosome positioning was found immediately upstream the stop codon, directly followed by a strong nucleosome valley in the case of Lml or a general decrease in signal in Lmb (Supplementary Figure 9A and 9B). In *B. cinerea* and *F. graminearum*, nucleosome signal was more defined with clear oscillations decaying past the stop codon (Supplementary Figures 9C and 9D). In *F. graminearum*, a lesser signal intensity seemed to characterise highly to moderately expressed genes (Supplementary Figure 9D), a feature visible only for highly expressed genes in *B. cinerea* (Supplementary Figure 9C). Altogether, our observations highlight the conservation of a nucleosome immediately before the stop codon, followed by a decrease in signal as a mark of gene expression in the four studied fungi. Markedly, when TTS of *B. cinerea* were considered, not expressed genes revealed a remarkably regular signal of weak amplitude. On the contrary, nucleosome signal around *F. graminearum* TTS of genes not expressed were characterized by a strong wave signal of well-arrayed nucleosomes (Supplementary Figure 9F). Recently, RSC was found to also play a positive effect on transcription termination, in addition to initiation (Ocampo *et al*., 2019). Considering that residence time of RNA polymerase II at the NDR downstream the TTS facilitates fast re-initiation of transcription (by recycling the RNA pol II; (Shandilya and Roberts, 2012; Cole *et al*., 2014), a scenario could be that RSC may be a transcription tuning knob, its presence stimulating transcription initiation while promoting RNA pol II dissociation to terminate transcription. Accordingly, the signal fuzziness we observed around the stop codons and TTS may reflect the highly dynamic nature of nucleosome positioning/removal catalyzed by RSC to regulate transcription termination. Overall, our results showed strong association of nucleosome landscapes at gene unit boundaries with expression levels in the four studied ascomycetes.

## CONCLUSION

The present study explored nucleosome landscapes of four phytopathogenic filamentous fungi with the aim of unravelling common features as well as potential specificities. Our general observation of nucleosome positioning genome-wide revealed shorter nucleosome-repeat lengths in *L. maculans* ‘brassicae’ and *L. maculans* ‘lepidii’ compared to *B. cinerea* and *F. graminearum*, suggesting a more compact chromatin fibre. High nucleosome occupancy was further observed in AT-rich regions of *L. maculans* ‘brassicae’, a feature *a priori* unexpected considering the well-described destabilising effect of AT-stretches but in line with the heterochromatic nature of these peculiar regions. High nucleosome occupancy was also observed at the *loci* of BOTY retrotransposons in the genome of *B. cinerea*. On the contrary, regions with reduced occupancy were observed in *F. graminearum* and co-localised with highly polymorphic regions described as prone to genetic evolution. As a whole, our results plead in favour of evolution of not only the positions of nucleosomes but also their occupancy, both likely hard-wired to genome sequence evolution, with regions defined by higher occupancies possibly securing sequence conservation. Evolution of genome sequences marked with peculiar chromatin signature profiles in relation with host adaptation has been previously described in the fungal pathogen *V. dahliae* (Cook *et al*., 2020). Considering how gene expression may relate on nucleosome patterning, an element of fungal specialization may rely on how chromatin remodeling proteins as well as promoter and other underlying genomic sequences have diversified.

## Data accessibility

All sequenced reads have been deposited with the Short Read Archive (SRA; https://www.ncbi.nlm.nih.gov/sra) under project accession number PRJNA580372. RNA-Seq data have been deposited in the Gene Expression Omnibus Database (GEO) (http://www.ncbi.nlm.nih.gov/geo/) under the entries GSE150127, GSE162838, and GSE162839.

## Availability

The MSTS (MNase-seq Tool Suite) is an open-source collection of tools developed by the BioinfoBIOGER platform by N. Lapalu and A. Simon, and available in the GitHub repository (https://github.com/nlapalu/MSTS).

## Acknowledgements

Version 3 of this preprint has been peer-reviewed and recommended by Peer Community In Genomics (https://doi.org/10.24072/pci.genomics.100014).

Sequencing was performed by the GenomEast platform, a member of the “France Génomique” consortium (ANR-10-INBS-0009). We particularly thank B. Jost for his help with setting up this project. We are grateful to the Genotoul bioinformatics platform Toulouse Midi-Pyrenees for providing computing resources.

This work was financially supported by the Plant Health and Environment division of the French National Institute for Agricultural Research (TACTIC Project AAP 2014). Funding for open access charge: Plant Health and Environment department of the French National Institute for Agricultural Research. J. L. Soyer was funded by a “Contrat Jeune Scientifique” grant from INRAE. The BIOGER Unit benefits from the support of Saclay Plant Sciences-SPS (ANR-17-EUR-0007).

## Supplementary data

### SUPPLEMENTARY TABLES

**Supplementary Table 1.** Summary of sequencing and mapping metrics

| Species | Culture condition | Analysis type | Biological replicate | Number of raw read pairs | Number of high-quality reads pairs* | Percentage of read pairs aligned to reference genome** |
| --- | --- | --- | --- | --- | --- | --- |
| *Fusarium graminearum* | Liquid MS, 3 days | MNase-seq | Fg#1 | 38,25,6895 | 36,297,487 | 88.1 |
|  |  |  | Fg#2 | 38,122,290 | 36,271,479 | 86.6 |
|  |  |  | Fg#3 | 37,122,439 | 35,234,660 | 88.7 |
|  |  | mRNA-seq | Fg#1 | 34,215,941 | 31,728,079 | 98.3 |
|  |  |  | Fg#2 | 37,209,377 | 34,506,796 | 98.3 |
|  |  |  | Fg#3 | 34,108,125 | 31,674,167 | 92.7 |
| *Leptosphaeria maculans* 'brassicae' | Fries 7 days | MNase-seq | Lmb#1 | 54,617,195 | 52,061,453 | 83.3 |
|  |  |  | Lmb#2 | 60,172,292 | 57,137,679 | 83.8 |
|  |  |  | Lmb#3 | 52,577,396 | 49,579,029 | 84.6 |
|  |  | mRNA-seq | Lmb#1 | 47,088,366 | 43,637,526 | 95.5 |
|  |  |  | Lmb#2 | 46,486,817 | 43,749,842 | 95.7 |
|  |  |  | Lmb#3 | 51,131,206 | 48,088,521 | 95.7 |
| *Leptosphaeria maculans* 'lepidii' | Fries 7 days | MNase-seq | Lml#1 | 41,026,352 | 39,044,192 | 92.9 |
|  |  |  | Lml#2 | 54,531,542 | 51,203,357 | 90.5 |
|  |  |  | Lml#3 | 55,487,109 | 52,896,833 | 91.6 |
|  |  | mRNA-seq | Lml#1 | 52,095,284 | 43,637,526 | 96.6 |
|  |  |  | Lml#2 | 50,886,249 | 49,097,848 | 96.8 |
|  |  |  | Lml#3 | 50,800,169 | 49,014,722 | 96.7 |
| *Botrytis cinerea* | Solid MM, 2 days | MNase-seq | Bc#1 | 36,861,968 | 34,606,209 | 94.8 |
|  |  |  | Bc#2 | 38,675,606 | 36,199,325 | 92.2 |
|  |  |  | Bc#3 | 36,684,375 | 34,437,003 | 95.0 |
|  |  | mRNA-seq | Bc#1 | 31,479,569 | 29,140,587 | 97.4 |
|  |  |  | Bc#2 | 38,826,778 | 35,931,380 | 97.6 |
|  |  |  | Bc#3 | 35,761,014 | 33,138,116 | 97.3 |

Sequencing reads for MNase-seq were deposited at SRA under accession number PRJNA580372, and those for RNA-seq at GEO repository under accession numbers GSE150127, GSE162838, and GSE162839; \*post quality trimming (see Materials and methods); \*\*concordantly exactly one time

**Supplementary Table 2.**
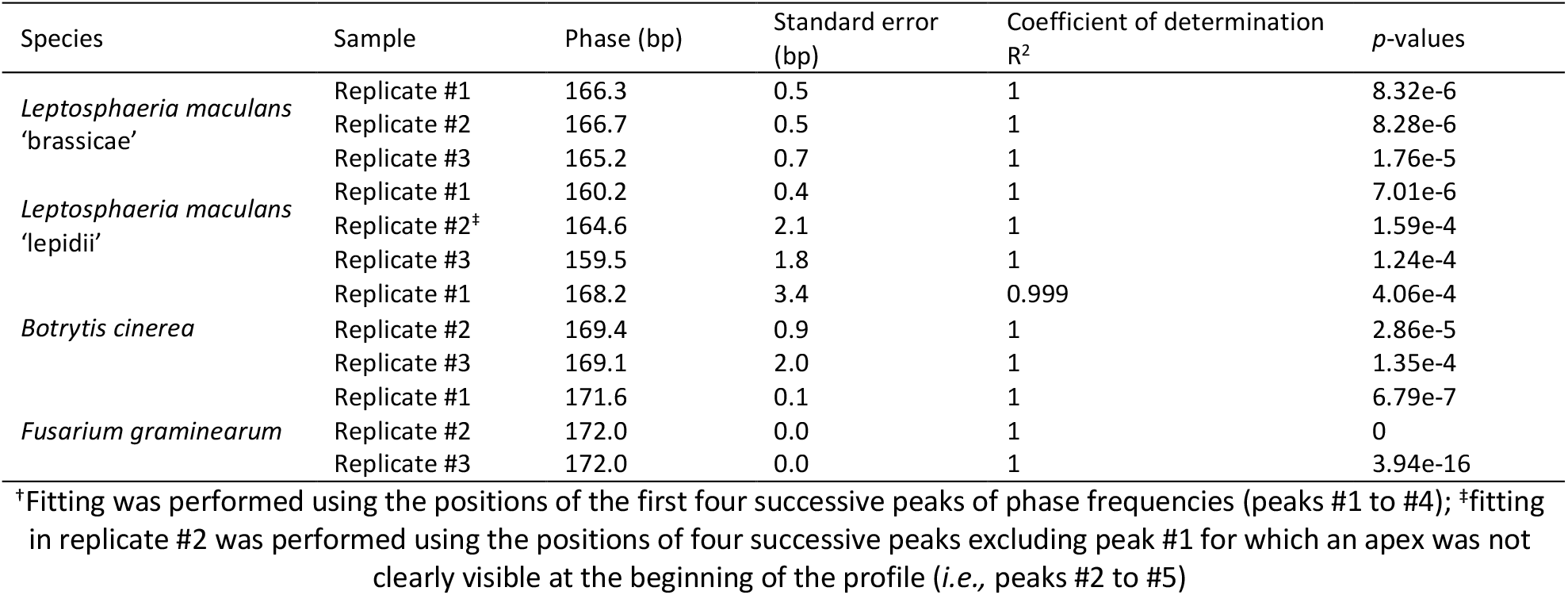
Summary of phases, standard errors, coefficient of determination R^2^, and *p*-values (*F*-test) determined after linear regression fitting^†^

**Supplementary Table 3.**
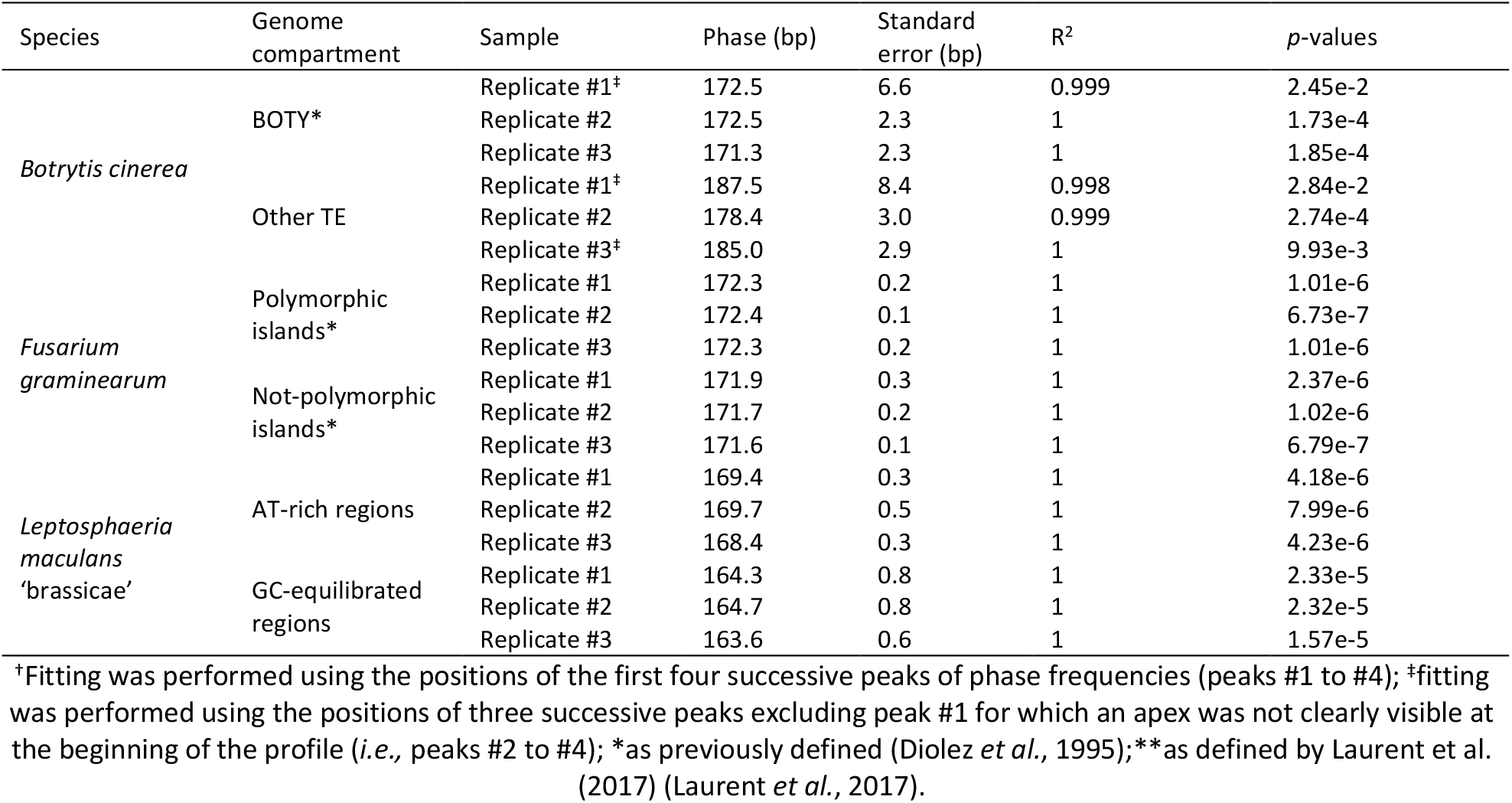
Summary of phases, standard errors, coefficient of determination R^2^, and *p*-values (*F*-test) determined after linear regression fitting^†^ in remarkable genome compartments for *Botrytis cinerea, Fusarium graminearum*, and *Leptosphaeria maculans* ‘brassicae’

**Supplementary Table 4.** Summary of periods, standard errors, coefficient of determination R^2^, and *p*-values (*F*-test) determined after linear regression fitting on autocorrelation plots for AA/AT/TT/TA and GG/GC/CC/CG frequencies

| Species | Di-nucleotides | Sample | Period (bp) | Standard error (bp) | $R^2$ | $p$-values |
| --- | --- | --- | --- | --- | --- | --- |
| *Leptosphaeria maculans* ‘brassicae’ | AA/AT/TT/TA | Replicate #1 | 9.8 | 0.1 | 0.999 | 3.67e-14 |
|  |  | Replicate #2 | 9.8 | 0.1 | 0.999 | 3.67e-14 |
|  |  | Replicate #3 | 9.7 | 0.1 | 0.999 | 1.64e-13 |
|  | GG/GC/CC/CG | Replicate #1 | 9.8 | 0.1 | 0.999 | 1.52e-13 |
|  |  | Replicate #2 | 9.7 | 0.1 | 0.999 | 3.36e-13 |
|  |  | Replicate #3 | 9.8 | 0.1 | 0.999 | 1.47e-13 |
| *Leptosphaeria maculans* ‘lepidii’ | AA/AT/TT/TA | Replicate #1 | 9.9 | 0.1 | 0.999 | 9.34e-13 |
|  |  | Replicate #2 | 9.8 | 0.1 | 0.999 | 1.47e-13 |
|  |  | Replicate #3 | 19.5 | 10.1 | 0.788 | 0.304 |
|  | GG/GC/CC/CG | Replicate #1 | 13.3 | 1 | 0.979 | 1.69e-4 |
|  |  | Replicate #2 | 11.3 | 0.5 | 0.989 | 4.2e-8 |
|  |  | Replicate #3 | 42 | NaN | NaN | NaN |
| *Botrytis cinerea* | AA/AT/TT/TA | Replicate #1 | 11.9 | 0.6 | 0.988 | 5.12e-6 |
|  |  | Replicate #2 | 11.9 | 0.3 | 0.996 | 2.32e-8 |
|  |  | Replicate #3 | 10.1 | 0.2 | 0.997 | 3.34e-10 |
|  | GG/GC/CC/CG | Replicate #1 | 14.3 | 1.9 | 0.949 | 4.91e-3 |
|  |  | Replicate #2 | 11.7 | 0.2 | 0.998 | 3.32e-9 |
|  |  | Replicate #3 | 11.2 | 0.3 | 0.995 | 3.52e-9 |
| *Fusarium graminearum* | AA/AT/TT/TA | Replicate #1 | 9.9 | 0.1 | 1 | 1.46e-14 |
|  |  | Replicate #2 | 9.9 | 0.1 | 1 | 1.46e-14 |
|  |  | Replicate #3 | 9.8 | 0.1 | 0.999 | 7.06e-14 |
|  | GG/GC/CC/CG | Replicate #1 | 9.8 | 0.1 | 0.999 | 2.08e-14 |
|  |  | Replicate #2 | 9.8 | 0.1 | 0.999 | 8.09e-14 |
|  |  | Replicate #3 | 9.8 | 0.1 | 0.999 | 4.96e-14 |

**Supplementary Table 5.** Summary of periods, standard errors, coefficient of determination R^2^, and *p*-values (*F*-test) determined after linear regression fitting on autocorrelation plots for AA/AT/TT/TA and GG/GC/CC/CG frequencies in AT-rich vs. GC-equilibrated regions of *Leptosphaeria maculans* ‘brassicae’

| Region | Di-nucleotides | Sample | Period (bp) | Standard error (bp) | $R^2$ | $p$-values |
| --- | --- | --- | --- | --- | --- | --- |
| AT-rich | AA/AT/TT/TA | Replicate #1 | 9.8 | 0.1 | 0.999 | 6.35e-14 |
|  |  | Replicate #2 | 9.9 | 0.1 | 0.999 | 2.4e-13 |
|  |  | Replicate #3 | 9.8 | 0.1 | 0.999 | 6.35e-14 |
|  | GG/GC/CC/CG | Replicate #1 | 9.8 | 0.1 | 0.999 | 2.27e-13 |
|  |  | Replicate #2 | 9.8 | 0.1 | 0.999 | 2.27e-13 |
|  |  | Replicate #3 | 9.8 | 0.1 | 0.999 | 8.09e-14 |
| GC-equilibrated | AA/AT/TT/TA | Replicate #1 | 9.8 | 0.1 | 0.999 | 5.63e-13 |
|  |  | Replicate #2 | 11.2 | 0.4 | 0.992 | 1.33e-8 |
|  |  | Replicate #3 | 9.7 | 0.1 | 0.999 | 3.36e-13 |
|  | GG/GC/CC/CG | Replicate #1 | 11.3 | 0.6 | 0.985 | 9.7e-7 |
|  |  | Replicate #2 | 9.8 | 0.2 | 0.996 | 6.34e-11 |
|  |  | Replicate #3 | 10.4 | 0.4 | 0.989 | 4.69e-8 |

**Supplementary Table 6.**
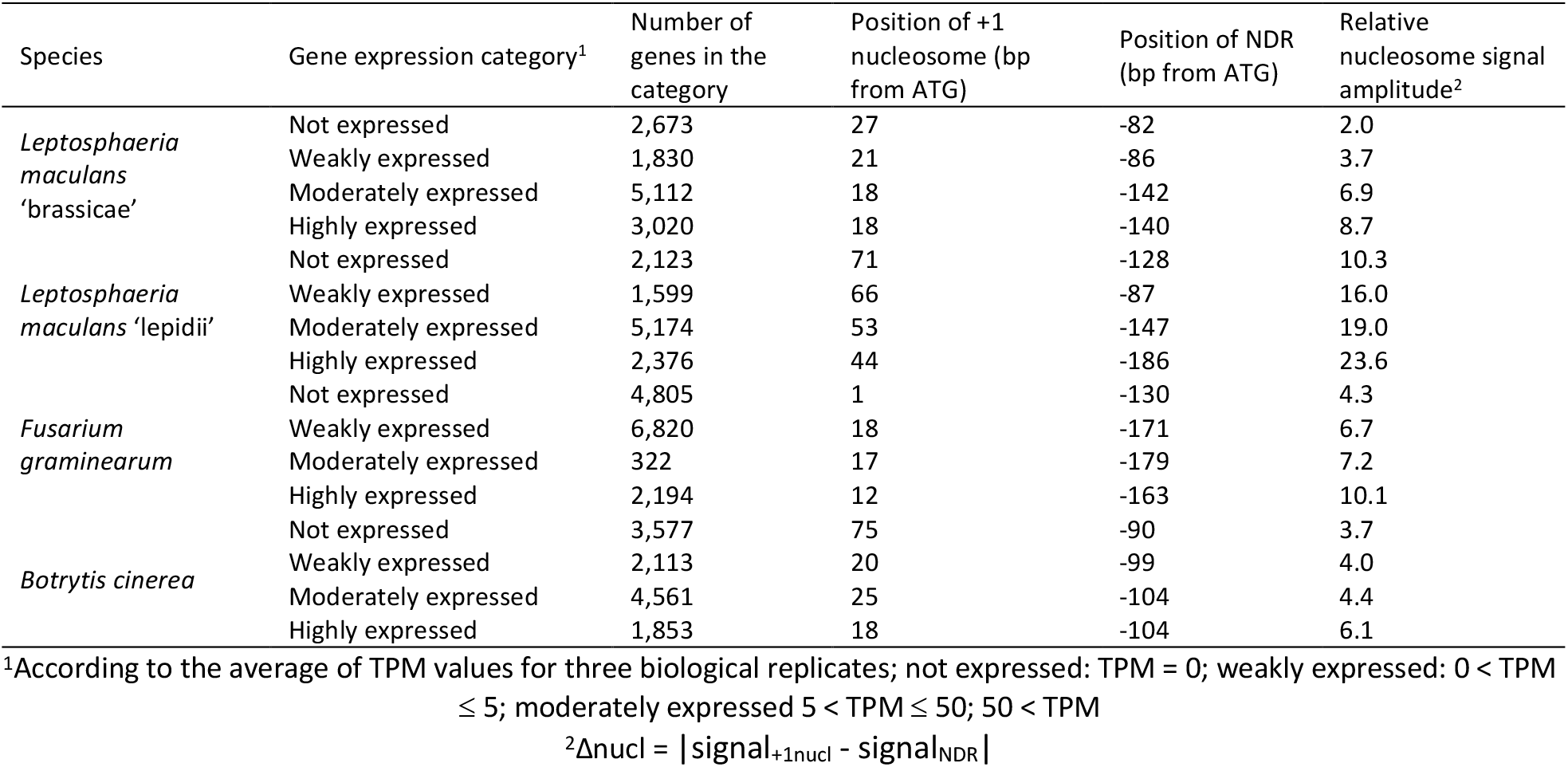
Nucleosome landscapes as a function of gene expression defining metrics

### SUPPLEMENTARY FIGURES

**Supplementary Figure 1.**
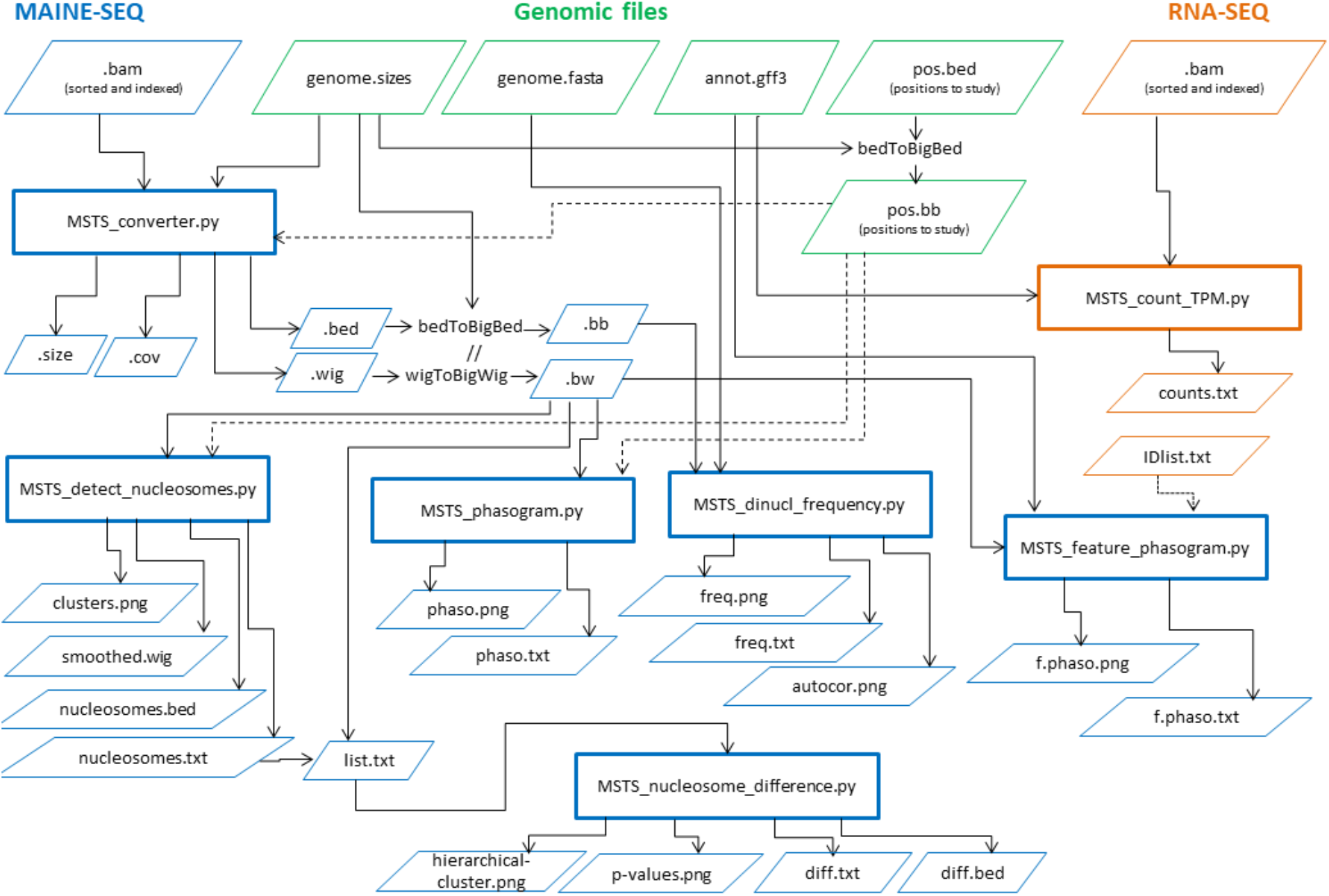
Workflow of the MNase-Seq Tool Suite (MSTS). The MSTS tool allows combining MNase-seq and RNA-seq data analyses, perform simple statistics and plot results. Detailed information can be found at https://github.com/nlapalu/MSTS.

**Supplementary Figure 2.**
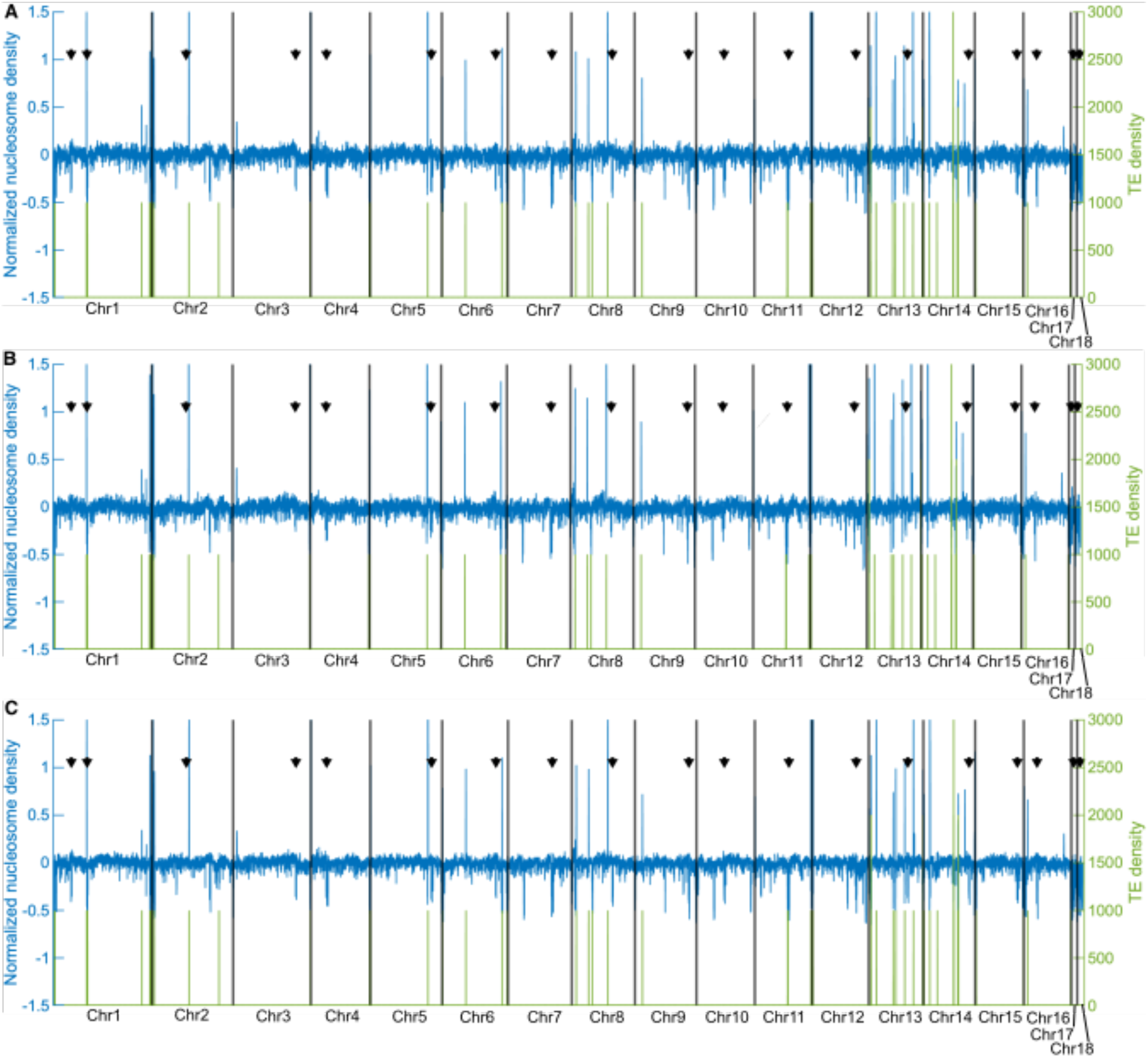
Nucleosome *vs*. BOTY TE density profiles in individual biological replicates of *Botrytis cinerea*. Coverage density profiles were computed for non-overlapping 1 kb-long bins along the chromosomes. In green are plotted BOTY transposable elements (TE) density profiles as previously described (Diolez *et al*., 1995). In blue are plotted the z-scored nucleosome density profiles. Black arrows indicate centromeres. (**A)**. Biological replicate #1; (**B)**. Biological Replicate #2; (**C)**. Biological replicate #3.

**Supplementary Figure 3.**
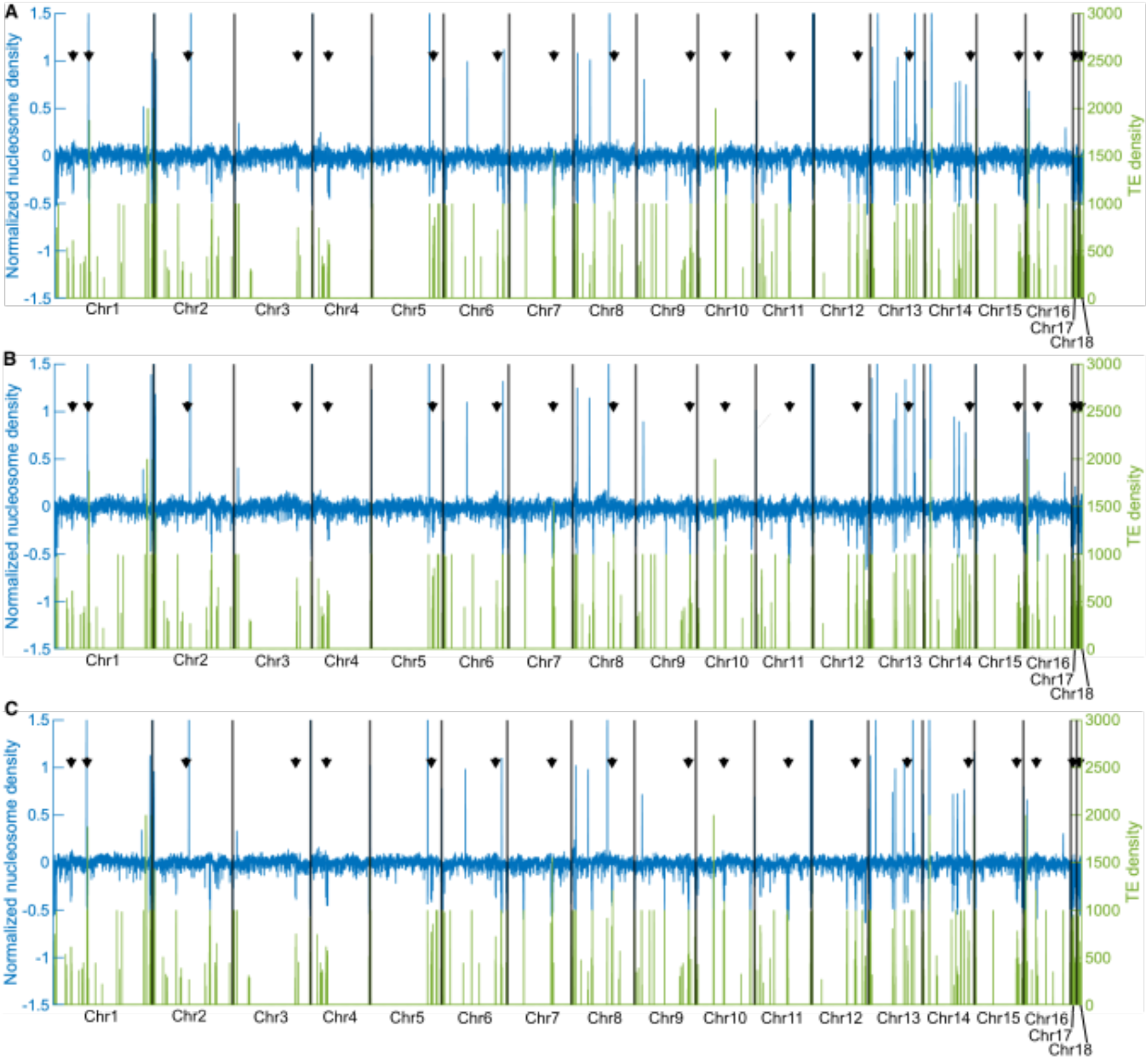
Nucleosome *vs*. TE (other than BOTY) density profiles in individual biological replicates of *Botrytis cinerea*. Coverage density profiles were computed for non-overlapping 1 kb-long bins along the chromosomes. In green are plotted TE (excluding BOTY) density profiles as previously described (Diolez *et al*., 1995). In blue are plotted the z-scored nucleosome density profiles. Black arrows indicate centromeres. (**A)**. Biological replicate #1; (**B)**. Biological Replicate #2; (**C)**. Biological replicate #3.

**Supplementary Figure 4.**
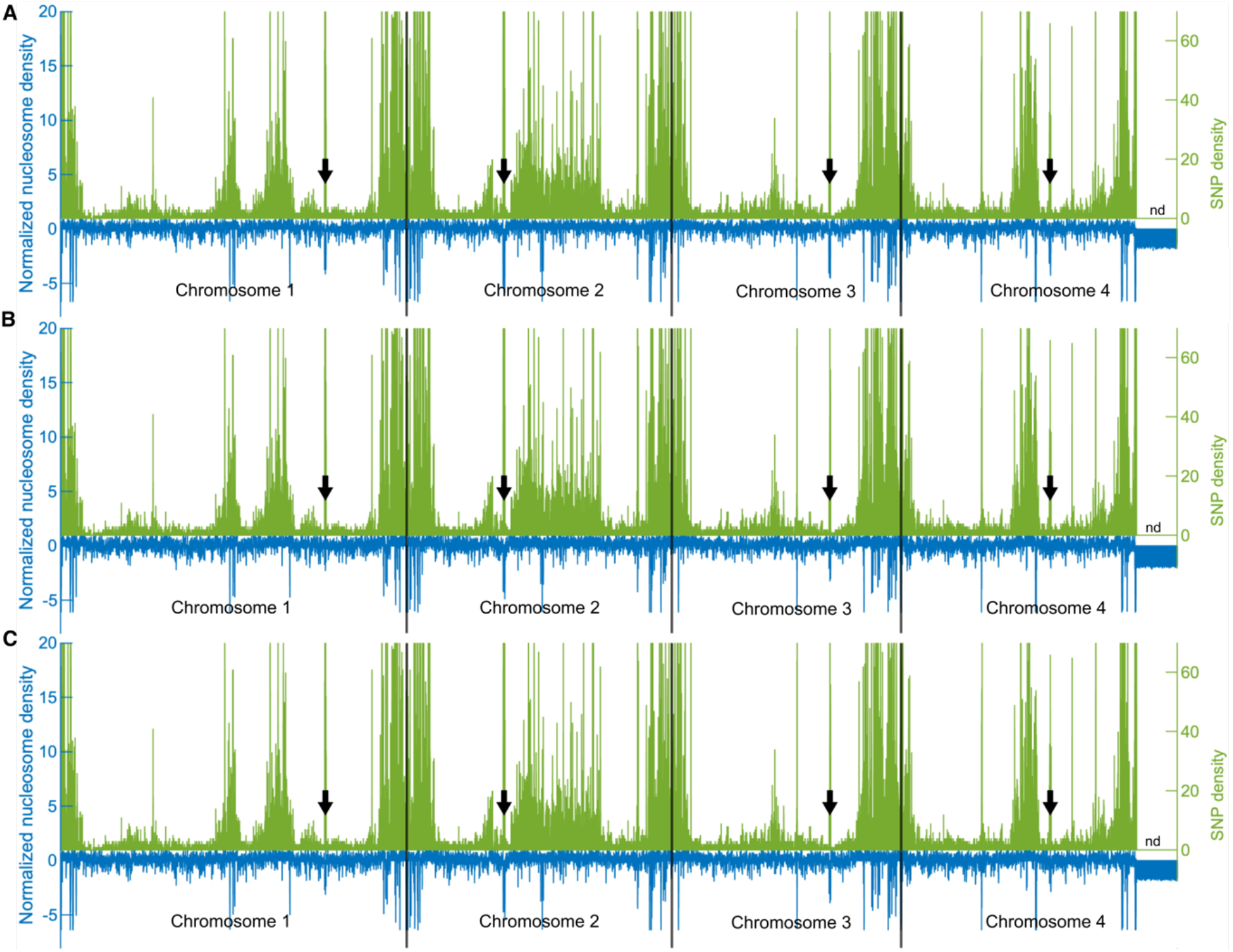
Nucleosome *vs*. SNP density profiles in individual biological replicates of *Fusarium graminearum*. Coverage density profiles were computed for non-overlapping 1 kb-long bins along the four chromosomes of *F. graminearum*. In green are plotted SNP density profiles as previously described (Laurent *et al*., 2017). “nd” indicates the highly variable 3’ end of chromosome 4 for which SNP were not called. In blue are plotted the z-scored nucleosome density profiles. Black arrows indicate centromeres (King *et al*., 2015); (**A)**. Biological replicate #1; (**B)**. Biological Replicate #2; (**C)**. Biological replicate #3.

**Supplementary Figure 5.**
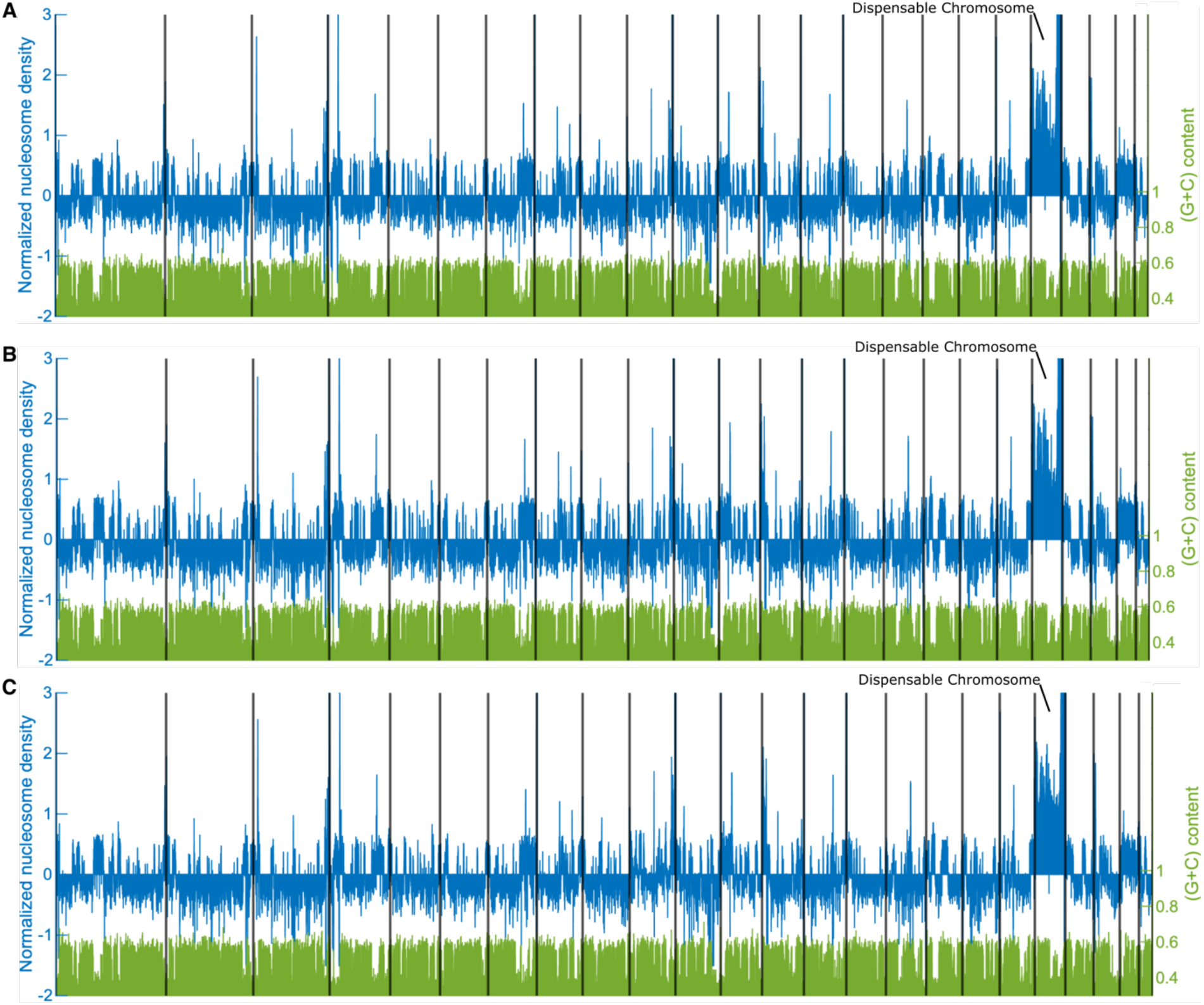
Normalized nucleosome density profiles *vs*. AT content in individual biological replicates of *Leptosphaeria maculans* ‘brassicae’. Z-scored coverage density profiles were computed for non-overlapping 1 kb-long bins along all supercontigs, separated by black lines. AT-rich 1 kb-long bins are plotted in green, isochores showing as dense regions (Rouxel *et al*., 2011). In blue are plotted the z-scored average nucleosome density profile (**A)**. Biological replicate #1; (**B)**. Biological Replicate #2; (**C)**. Biological replicate #3.

**Supplementary Figure 6.**
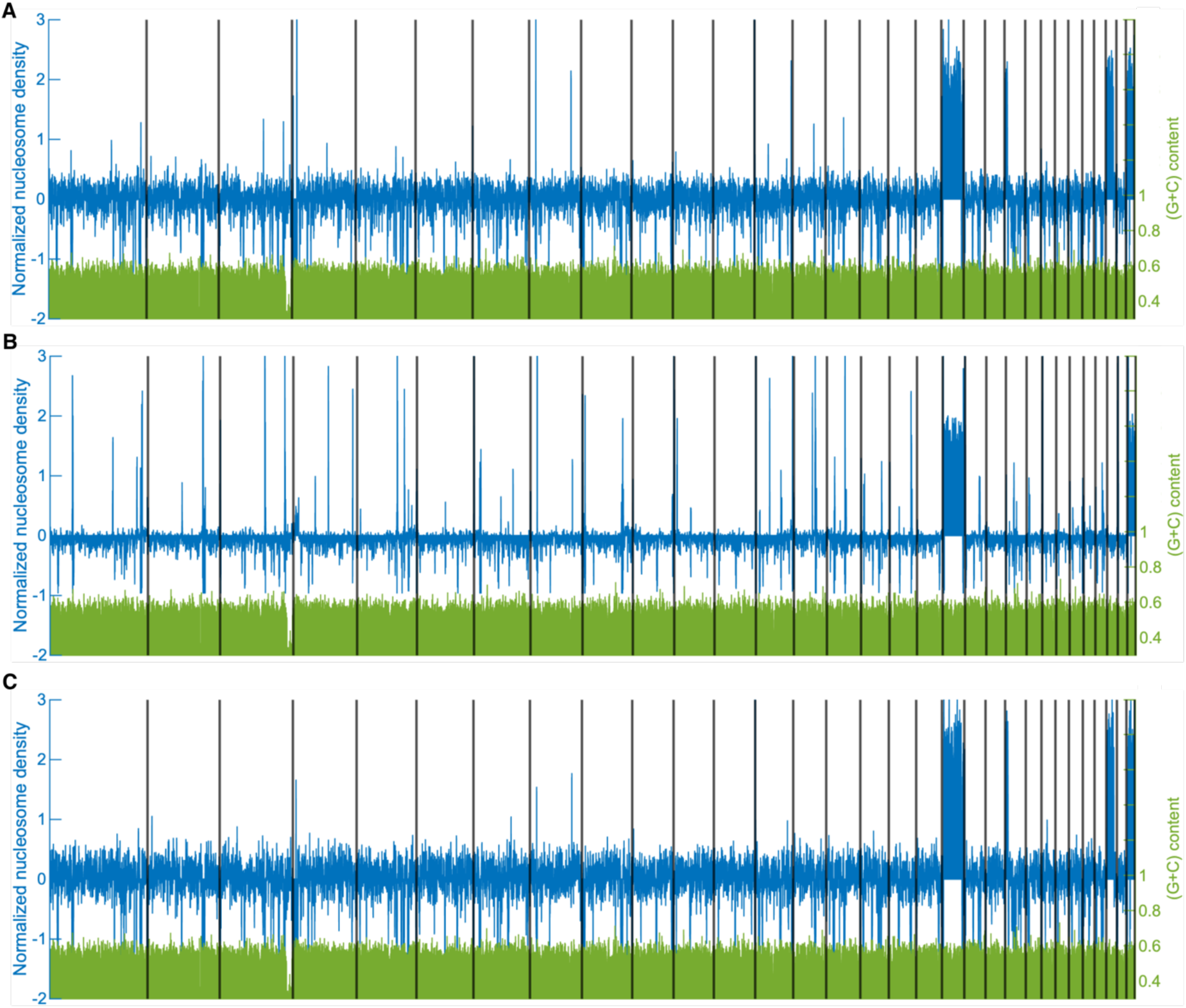
Normalized nucleosome density profiles *vs*. AT content in individual biological replicates of *Leptosphaeria maculans* ‘lepidii’. Coverage density profiles were computed for non-overlapping 1 kb-long bins along all supercontigs, separated by black lines. AT-rich 1 kb-long bins are plotted in green, with no isochore visible. In blue are plotted the z-scored average nucleosome density profile (**A)**. Biological replicate #1; (**B)**. Biological Replicate #2; (**C)**. Biological replicate #3.

**Supplementary Figure 7.**
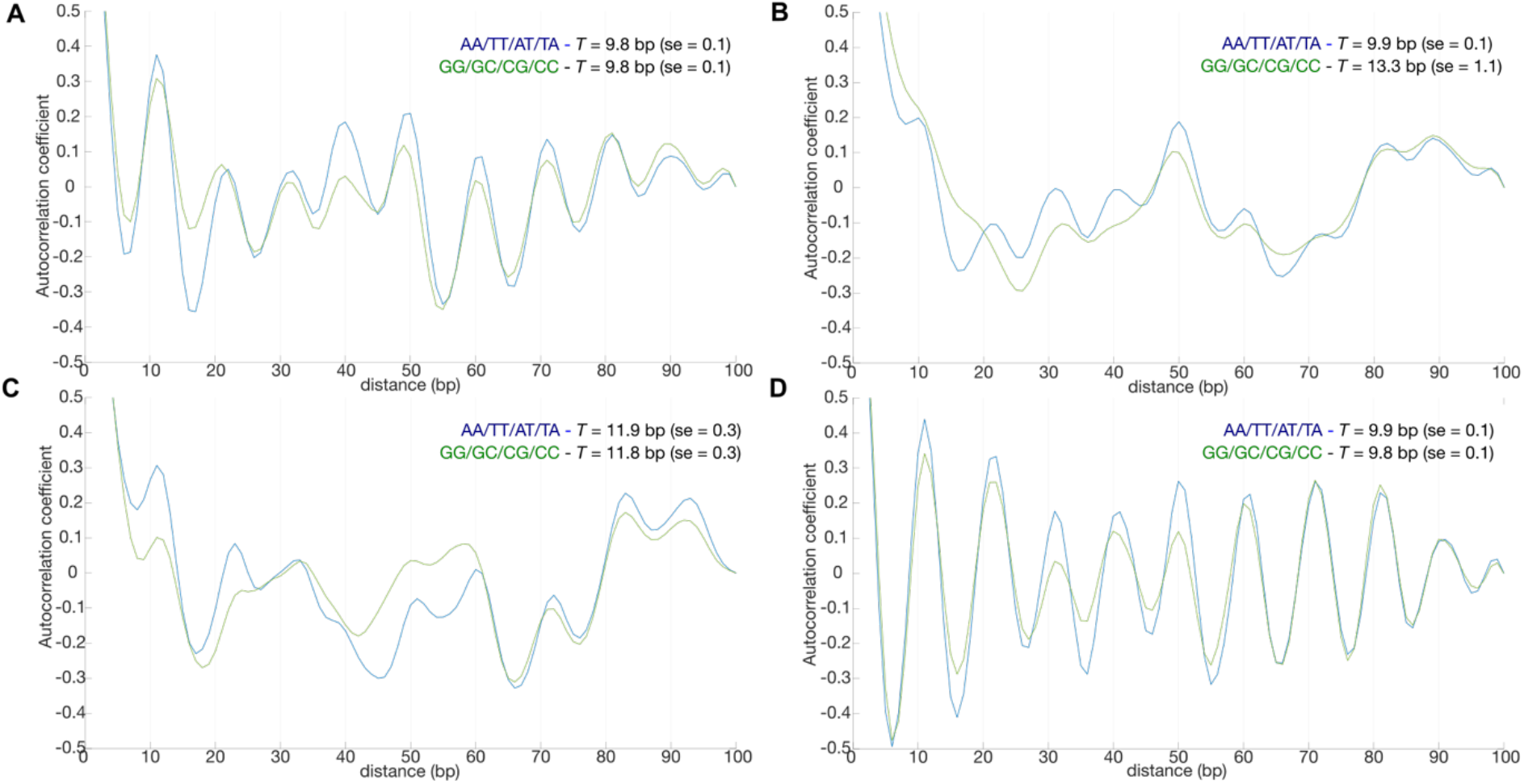
Autocorrelation coefficients of di-nucleotide frequencies (average of three biological replicates) for *Leptosphaeria maculans* ‘brassicae’ (**A**), L*eptosphaeria maculans* ‘lepidii’ (**B**), *Botrytis cinerea* (**C**), and *Fusarium graminearum* (**D**). In blue: autocorrelation for AA/TT/AT/TA frequencies; in green: autocorrelation for GG/CC/CG/GC frequencies. Periodicities *T* were inferred after linear regression fitting to peak positions as a function of peak rank. Periods, standard errors (se), R^2^ (coefficient of determination), and *p*-values (*F*-test) were also determined individually for each replicate (see Supplementary Table 4).

**Supplementary Figure 8.**
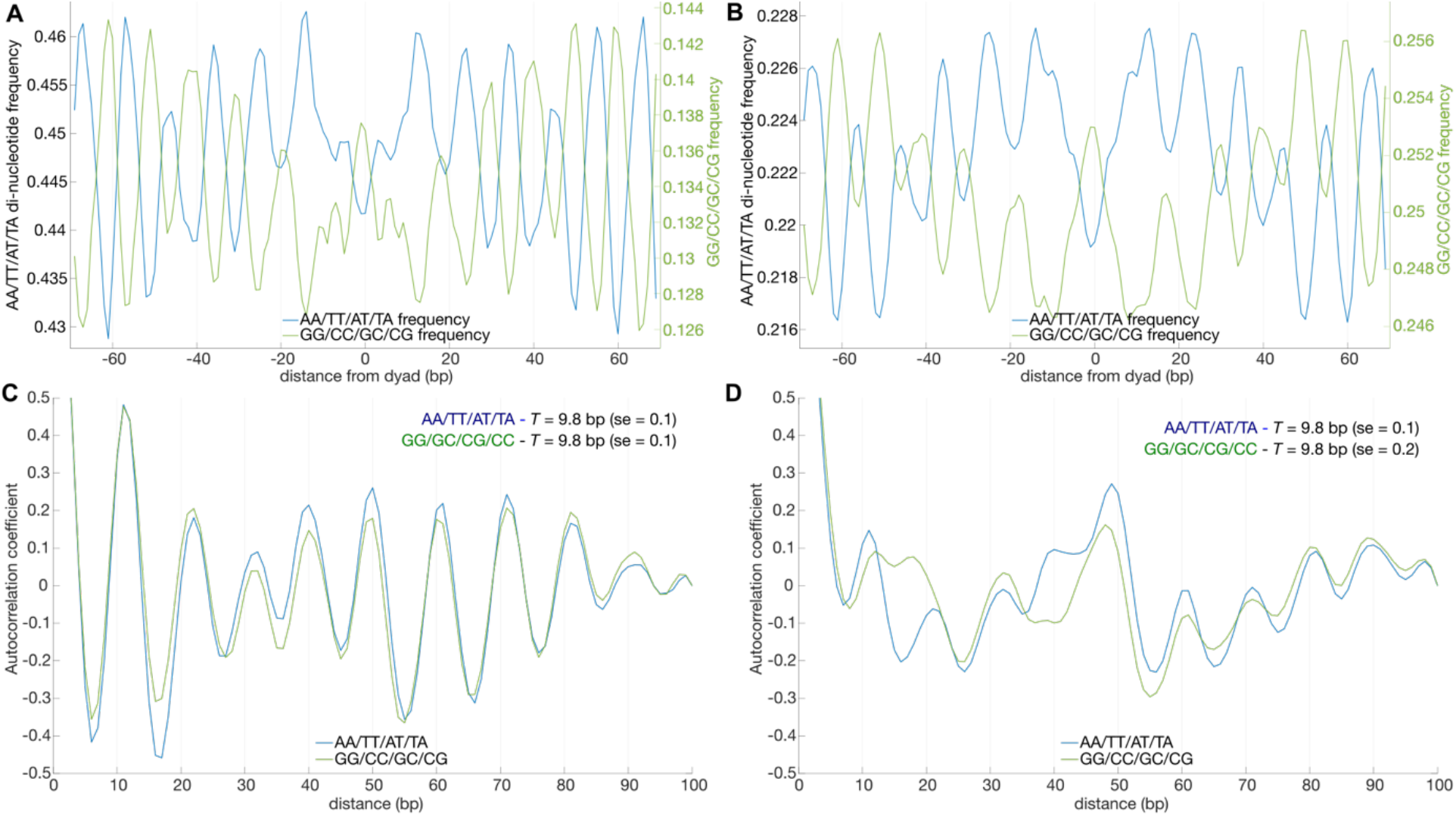
Repeated di-nucleotide patterns in nucleosomal DNA located in AT-rich and GC-equilibrated regions of *Leptosphaeria maculans* ‘brassicae’. **A** and **B**. Normalized di-nucleotides frequency plots (average of three biological replicates) for nucleosomal DNA in AT-rich regions (**A**) and GC-equilibrated regions (**B**). **C** and **D**. Autocorrelation coefficients of di-nucleotides frequencies (average of three biological replicates) for nucleosomal DNA in AT-rich regions (**C**) and GC-equilibrated regions (**D**).

**Supplementary Figure 9.**
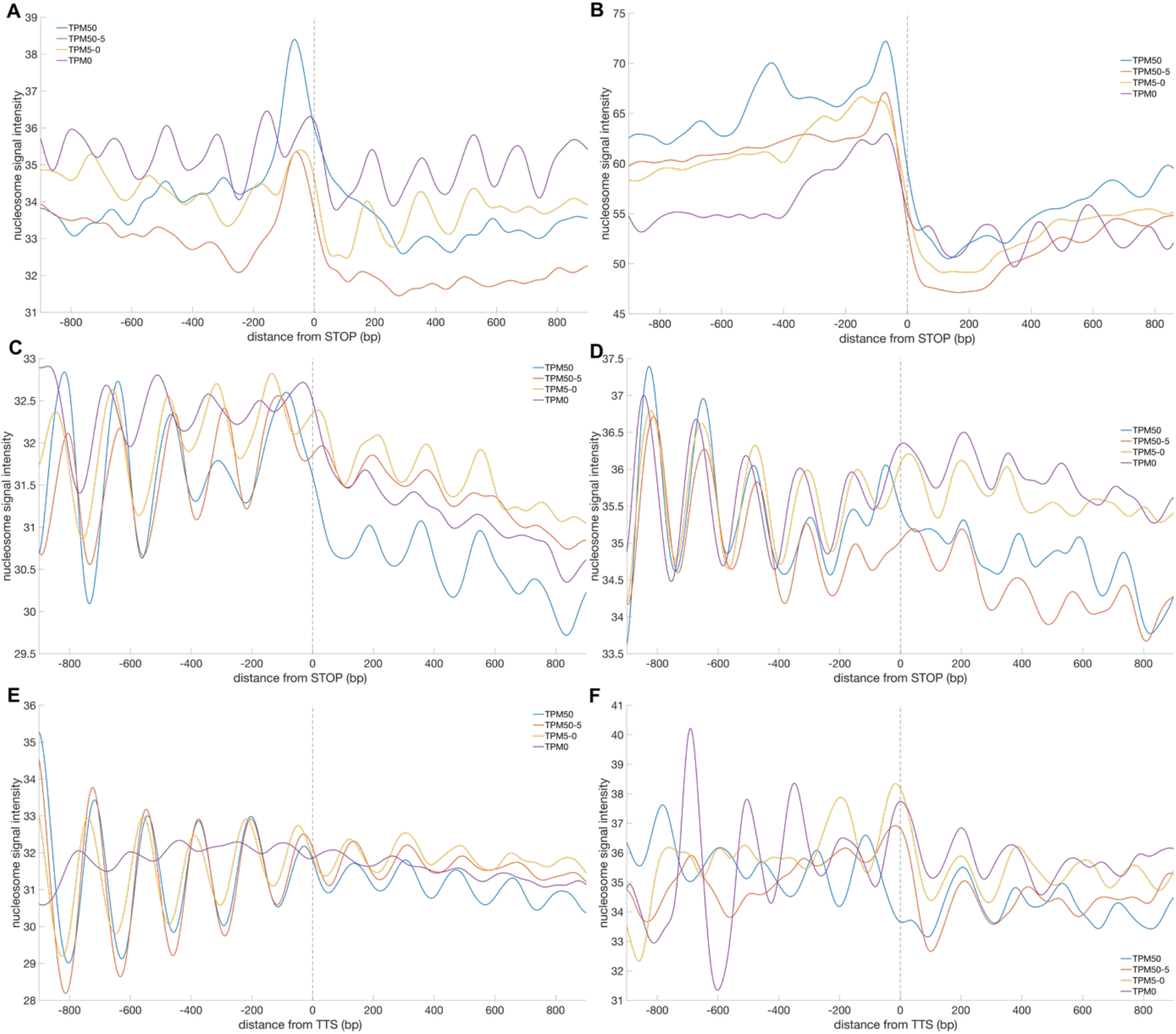
Nucleosome organisation at stop codons/TTS *vs*. gene expression. Average (three biological replicates for each fungus/condition) nucleosome signal as a function of position (in base pairs) relative to the stop codon (and TTS for *F. graminearum* and *B. cinerea*). TPM = Transcripts Per Million. A. In Lmb, Stop-centred; B. In Lml, Stop-centred; C. In *B. cinerea*, Stop-centred; D. *F. graminearum*, Stop-centred; E. In *B. cinerea*, TTS-centred; F. *F. graminearum*, TTS-centred.

## References

Amselem, J. et al. (2011) ‘Genomic analysis of the necrotrophic fungal pathogens Sclerotinia sclerotiorum and Botrytis cinerea’, PLoS genetics, 7(8), p. e1002230. https://doi.org/10.1371/journal.pgen.1002230.

Balesdent, M.H. et al. (2001) ‘Genetic Control and Host Range of Avirulence Toward Brassica napus Cultivars Quinta and Jet Neuf in Leptosphaeria maculans’, Phytopathology, 91(1), pp. 70–76. https://doi.org/10.1094/PHYTO.2001.91.1.70.

Barnes, T. and Korber, P. (2021) ‘The Active Mechanism of Nucleosome Depletion by Poly(dA:dT) Tracts In Vivo’, International Journal of Molecular Sciences, 22(15), p. 8233. https://doi.org/10.3390/ijms22158233.

Basenko, E.Y. et al. (2018) ‘FungiDB: An Integrated Bioinformatic Resource for Fungi and Oomycetes’, Journal of Fungi (Basel, Switzerland), 4(1). https://doi.org/10.3390/jof4010039.

Bolger, A.M., Lohse, M. and Usadel, B. (2014) ‘Trimmomatic: a flexible trimmer for Illumina sequence data’, Bioinformatics (Oxford, England), 30(15), pp. 2114–2120. https://doi.org/10.1093/bioinformatics/btu170.

Boutigny, A.-L. et al. (2009) ‘Ferulic acid, an efficient inhibitor of type B trichothecene biosynthesis and Tri gene expression in Fusarium liquid cultures’, Mycological Research, 113(Pt 6-7), pp. 746–753. https://doi.org/10.1016/j.mycres.2009.02.010.

Bunnik, E.M. et al. (2014) ‘DNA-encoded nucleosome occupancy is associated with transcription levels in the human malaria parasite Plasmodium falciparum’, BMC genomics, 15, p. 347. https://doi.org/10.1186/1471-2164-15-347.

Chen, K. et al. (2013) ‘DANPOS: Dynamic analysis of nucleosome position and occupancy by sequencing’, Genome Research, 23(2), pp. 341–351. https://doi.org/10.1101/gr.142067.112.

Cole, H.A. et al. (2014) ‘Heavy transcription of yeast genes correlates with differential loss of histone H2B relative to H4 and queued RNA polymerases’, Nucleic Acids Research, 42(20), pp. 12512–12522. https://doi.org/10.1093/nar/gku1013.

Collemare, J., O’Connell, R. and Lebrun, M.-H. (2019) ‘Nonproteinaceous effectors: the terra incognita of plant-fungal interactions’, The New Phytologist, 223(2), pp. 590–596. https://doi.org/10.1111/nph.15785.

Collemare, J. and Seidl, M.F. (2019) ‘Chromatin-dependent regulation of secondary metabolite biosynthesis in fungi: is the picture complete?’, FEMS microbiology reviews, 43(6), pp. 591–607. https://doi.org/10.1093/femsre/fuz018.

Cook, D.E. et al. (2020) ‘A unique chromatin profile defines adaptive genomic regions in a fungal plant pathogen’, eLife, 9, p. e62208. https://doi.org/10.7554/eLife.62208.

Core, L.J., Waterfall, J.J. and Lis, J.T. (2008) ‘Nascent RNA sequencing reveals widespread pausing and divergent initiation at human promoters’, Science (New York, N.Y.), 322(5909), pp. 1845–1848. https://doi.org/10.1126/science.1162228.

Cuomo, C.A. et al. (2007) ‘The Fusarium graminearum genome reveals a link between localized polymorphism and pathogen specialization’, Science (New York, N.Y.), 317(5843), pp. 1400–1402. https://doi.org/10.1126/science.1143708.

Ding, Y., Gardiner, D.M. and Kazan, K. (2022) ‘Transcriptome analysis reveals infection strategies employed by Fusarium graminearum as a root pathogen’, Microbiological Research, 256, p. 126951. https://doi.org/10.1016/j.micres.2021.126951.

Dobin, A. et al. (2013) ‘STAR: ultrafast universal RNA-seq aligner’, Bioinformatics (Oxford, England), 29(1), pp. 15–21. https://doi.org/10.1093/bioinformatics/bts635.

Dutreux, F. et al. (2018) ‘De novo assembly and annotation of three Leptosphaeria genomes using Oxford Nanopore MinION sequencing’, Scientific Data, 5, p. 180235. https://doi.org/10.1038/sdata.2018.235.

Fisher, M.C. et al. (2012) ‘Emerging fungal threats to animal, plant and ecosystem health’, Nature, 484(7393), pp. 186–194. https://doi.org/10.1038/nature10947.

Flores, O. and Orozco, M. (2011) ‘nucleR: a package for non-parametric nucleosome positioning’, Bioinformatics, 27(15), pp. 2149–2150. https://doi.org/10.1093/bioinformatics/btr345.

Fudal, I. et al. (2007) ‘Heterochromatin-like regions as ecological niches for avirulence genes in the Leptosphaeria maculans genome: map-based cloning of AvrLm6’, Molecular plant-microbe interactions: MPMI, 20(4), pp. 459–470. https://doi.org/10.1094/MPMI-20-4-0459.

Gay, E.J. et al. (2020) ‘Large-scale transcriptomics to dissect two years of the life of a fungal phytopathogen interacting with its host plant’, bioRxiv, p. 2020.10.13.331520. https://doi.org/10.1101/2020.10.13.331520.

Ghassabi Kondalaji, S. and Bowman, G.D. (2021) ‘Reb1, Cbf1, and Pho4 bias histone sliding and deposition away from their binding sites’, Molecular and Cellular Biology, p. MCB0047221. https://doi.org/10.1128/MCB.00472-21.

Grandaubert, J. et al. (2014) ‘Transposable element-assisted evolution and adaptation to host plant within the Leptosphaeria maculans-Leptosphaeria biglobosa species complex of fungal pathogens’, BMC genomics, 15, p. 891. https://doi.org/10.1186/1471-2164-15-891.

Hallen, H.E. et al. (2007) ‘Gene expression shifts during perithecium development in Gibberella zeae (anamorph Fusarium graminearum), with particular emphasis on ion transport proteins’, Fungal genetics and biology: FG & B, 44(11), pp. 1146–1156. https://doi.org/10.1016/j.fgb.2007.04.007.

Hawksworth, D.L. and Lücking, R. (2017) ‘Fungal Diversity Revisited: 2.2 to 3.8 Million Species’, Microbiology Spectrum, 5(4). https://doi.org/10.1128/microbiolspec.FUNK-0052-2016.

Hornung, G., Oren, M. and Barkai, N. (2012) ‘Nucleosome organization affects the sensitivity of gene expression to promoter mutations’, Molecular Cell, 46(3), pp. 362–368. https://doi.org/10.1016/j.molcel.2012.02.019.

Hu, S. et al. (2017) ‘CAM: A quality control pipeline for MNase-seq data’, PLOS ONE, 12(8), p. e0182771. https://doi.org/10.1371/journal.pone.0182771.

Hughes, A.L. and Rando, O.J. (2015) ‘Comparative Genomics Reveals Chd1 as a Determinant of Nucleosome Spacing in Vivo’, G3 (Bethesda, Md.), 5(9), pp. 1889–1897. https://doi.org/10.1534/g3.115.020271.

Jiang, C. and Pugh, B.F. (2009) ‘A compiled and systematic reference map of nucleosome positions across the Saccharomyces cerevisiae genome’, Genome Biology, 10(10), p. R109. https://doi.org/10.1186/gb-2009-10-10-r109.

Jin, H., Finnegan, A.I. and Song, J.S. (2018) ‘A unified computational framework for modeling genome-wide nucleosome landscape’, Physical Biology, 15(6), p. 066011. https://doi.org/10.1088/1478-3975/aadad2.

Kaplan, N. et al. (2009) ‘The DNA-encoded nucleosome organization of a eukaryotic genome’, Nature, 458(7236), pp. 362–366. https://doi.org/10.1038/nature07667.

Kelloniemi, J. et al. (2015) ‘Analysis of the Molecular Dialogue Between Gray Mold (Botrytis cinerea) and Grapevine (Vitis vinifera) Reveals a Clear Shift in Defense Mechanisms During Berry Ripening’, Molecular plant-microbe interactions: MPMI, 28(11), pp. 1167–1180. https://doi.org/10.1094/MPMI-02-15-0039-R.

King, R. et al. (2015) ‘The completed genome sequence of the pathogenic ascomycete fungus Fusarium graminearum’, BMC genomics, 16, p. 544. https://doi.org/10.1186/s12864-015-1756-1.

King, R., Urban, M. and Hammond-Kosack, K.E. (2017) ‘Annotation of Fusarium graminearum (PH-1) Version 5.0’, Genome Announcements, 5(2). https://doi.org/10.1128/genomeA.01479-16.

Krietenstein, N. et al. (2016) ‘Genomic Nucleosome Organization Reconstituted with Pure Proteins’, Cell, 167(3), pp. 709-721.e12. https://doi.org/10.1016/j.cell.2016.09.045.

Kwak, H. et al. (2013) ‘Precise maps of RNA polymerase reveal how promoters direct initiation and pausing’, Science (New York, N.Y.), 339(6122), pp. 950–953. https://doi.org/10.1126/science.1229386.

Langmead, B. and Salzberg, S.L. (2012) ‘Fast gapped-read alignment with Bowtie 2’, Nature Methods, 9(4), pp. 357–359. https://doi.org/10.1038/nmeth.1923.

Lantermann, A.B. et al. (2010) ‘Schizosaccharomyces pombe genome-wide nucleosome mapping reveals positioning mechanisms distinct from those of Saccharomyces cerevisiae’, Nature Structural & Molecular Biology, 17(2), pp. 251–257. https://doi.org/10.1038/nsmb.1741.

Laurent, B. et al. (2017) ‘Landscape of genomic diversity and host adaptation in Fusarium graminearum’, BMC genomics, 18(1), p. 203. https://doi.org/10.1186/s12864-017-3524-x.

Li, B. et al. (2010) ‘RNA-Seq gene expression estimation with read mapping uncertainty’, Bioinformatics, 26(4), pp. 493–500. https://doi.org/10.1093/bioinformatics/btp692.

Lieleg, C. et al. (2015) ‘Nucleosome spacing generated by ISWI and CHD1 remodelers is constant regardless of nucleosome density’, Molecular and Cellular Biology, 35(9), pp. 1588–1605. https://doi.org/10.1128/MCB.01070-14.

Lo Presti, L. et al. (2015) ‘Fungal effectors and plant susceptibility’, Annual Review of Plant Biology, 66, pp. 513–545. https://doi.org/10.1146/annurev-arplant-043014-114623.

Locke, G. et al. (2013) ‘Global remodeling of nucleosome positions in C. elegans’, BMC genomics, 14, p. 284. https://doi.org/10.1186/1471-2164-14-284.

Min, I.M. et al. (2011) ‘Regulating RNA polymerase pausing and transcription elongation in embryonic stem cells’, Genes & Development, 25(7), pp. 742–754. https://doi.org/10.1101/gad.2005511.

Nishida, H. et al. (2009) ‘Genome-wide maps of mono- and di-nucleosomes of Aspergillus fumigatus’, Bioinformatics (Oxford, England), 25(18), pp. 2295–2297. https://doi.org/10.1093/bioinformatics/btp413.

Nishida, H. et al. (2012) ‘Characteristics of nucleosomes and linker DNA regions on the genome of the basidiomycete Mixia osmundae revealed by mono- and dinucleosome mapping’, Open Biology, 2(4), p. 120043. https://doi.org/10.1098/rsob.120043.

Oberbeckmann, E., Krietenstein, N., et al. (2021) ‘Genome information processing by the INO80 chromatin remodeler positions nucleosomes’, Nature Communications, 12(1), p. 3231. https://doi.org/10.1038/s41467-021-23016-z.

Oberbeckmann, E., Niebauer, V., et al. (2021) ‘Ruler elements in chromatin remodelers set nucleosome array spacing and phasing’, Nature Communications, 12(1), p. 3232. https://doi.org/10.1038/s41467-021-23015-0.

Ocampo, J. et al. (2016) ‘The ISW1 and CHD1 ATP-dependent chromatin remodelers compete to set nucleosome spacing in vivo’, Nucleic Acids Research, 44(10), pp. 4625–4635. https://doi.org/10.1093/nar/gkw068.

Ocampo, J. et al. (2019) ‘Contrasting roles of the RSC and ISW1/CHD1 chromatin remodelers in RNA polymerase II elongation and termination’, Genome Research, 29(3), pp. 407–417. https://doi.org/10.1101/gr.242032.118.

Pedro, H. et al. (2019) ‘Collaborative Annotation Redefines Gene Sets for Crucial Phytopathogens’, Frontiers in Microbiology, 10, p. 2477. https://doi.org/10.3389/fmicb.2019.02477.

Ponts, N. et al. (2010) ‘Nucleosome landscape and control of transcription in the human malaria parasite’, Genome Research, 20(2), pp. 228–238. https://doi.org/10.1101/gr.101063.109.

Porquier, A. et al. (2016) ‘The botrydial biosynthetic gene cluster of Botrytis cinerea displays a bipartite genomic structure and is positively regulated by the putative Zn(II)2Cys6 transcription factor BcBot6’, Fungal genetics and biology: FG & B, 96, pp. 33–46. https://doi.org/10.1016/j.fgb.2016.10.003.

Porquier, A. et al. (2021) ‘Retrotransposons as pathogenicity factors of the plant pathogenic fungus Botrytis cinerea’, bioRxiv, p. 2021.04.13.439636. https://doi.org/10.1101/2021.04.13.439636.

Radman-Livaja, M. and Rando, O.J. (2010) ‘Nucleosome positioning: how is it established, and why does it matter?’, Developmental Biology, 339(2), pp. 258–266. https://doi.org/10.1016/j.ydbio.2009.06.012.

Reyes, A.A., Marcum, R.D. and He, Y. (2021) ‘Structure and Function of Chromatin Remodelers’, Journal of Molecular Biology, 433(14), p. 166929. https://doi.org/10.1016/j.jmb.2021.166929.

Richmond, T.J. and Davey, C.A. (2003) ‘The structure of DNA in the nucleosome core’, Nature, 423(6936), pp. 145–150. https://doi.org/10.1038/nature01595.

Robinson, P.J.J. et al. (2006) ‘EM measurements define the dimensions of the “30-nm” chromatin fiber: evidence for a compact, interdigitated structure’, Proceedings of the National Academy of Sciences of the United States of America, 103(17), pp. 6506–6511. https://doi.org/10.1073/pnas.0601212103.

Rouxel, T. et al. (2011) ‘Effector diversification within compartments of the Leptosphaeria maculans genome affected by Repeat-Induced Point mutations’, Nature Communications, 2, p. 202. https://doi.org/10.1038/ncomms1189.

Russell, K. et al. (2014) ‘Homopolymer tract organization in the human malarial parasite Plasmodium falciparum and related Apicomplexan parasites’, BMC genomics, 15, p. 848. https://doi.org/10.1186/1471-2164-15-848.

Sánchez-Vallet, A. et al. (2018) ‘The Genome Biology of Effector Gene Evolution in Filamentous Plant Pathogens’, Annual Review of Phytopathology, 56(1), pp. 21–40. https://doi.org/10.1146/annurev-phyto-080516-035303.

Satchwell, S.C., Drew, H.R. and Travers, A.A. (1986) ‘Sequence periodicities in chicken nucleosome core DNA’, Journal of Molecular Biology, 191(4), pp. 659–675. https://doi.org/10.1016/0022-2836(86)90452-3.

Segal, E. et al. (2006) ‘A genomic code for nucleosome positioning’, Nature, 442(7104), pp. 772–778. https://doi.org/10.1038/nature04979.

Segal, E. and Widom, J. (2009a) ‘Poly(dA:dT) tracts: major determinants of nucleosome organization’, Current Opinion in Structural Biology, 19(1), pp. 65–71. https://doi.org/10.1016/j.sbi.2009.01.004.

Segal, E. and Widom, J. (2009b) ‘What controls nucleosome positions?’, Trends in genetics: TIG, 25(8), pp. 335–343. https://doi.org/10.1016/j.tig.2009.06.002.

Shandilya, J. and Roberts, S.G.E. (2012) ‘The transcription cycle in eukaryotes: from productive initiation to RNA polymerase II recycling’, Biochimica Et Biophysica Acta, 1819(5), pp. 391–400. https://doi.org/10.1016/j.bbagrm.2012.01.010.

Simão, F.A. et al. (2015) ‘BUSCO: assessing genome assembly and annotation completeness with single-copy orthologs’, Bioinformatics, 31(19), pp. 3210–3212. https://doi.org/10.1093/bioinformatics/btv351.

Simon, A. et al. (2013) ‘Screening of a Botrytis cinerea one-hybrid library reveals a Cys2His2 transcription factor involved in the regulation of secondary metabolism gene clusters’, Fungal genetics and biology: FG & B, 52, pp. 9–19. https://doi.org/10.1016/j.fgb.2013.01.006.

Singh, A.K. and Mueller-Planitz, F. (2021) ‘Nucleosome Positioning and Spacing: From Mechanism to Function’, Journal of Molecular Biology, 433(6), p. 166847. https://doi.org/10.1016/j.jmb.2021.166847.

Soyer, J.L. et al. (2014) ‘Epigenetic control of effector gene expression in the plant pathogenic fungus Leptosphaeria maculans’, PLoS genetics, 10(3), p. e1004227. https://doi.org/10.1371/journal.pgen.1004227.

Soyer, J.L. et al. (2015) ‘Chromatin analyses of Zymoseptoria tritici: Methods for chromatin immunoprecipitation followed by high-throughput sequencing (ChIP-seq)’, Fungal genetics and biology: FG & B, 79, pp. 63–70. https://doi.org/10.1016/j.fgb.2015.03.006.

Soyer, J.L. et al. (2018) ‘To B or not to B: a tale of unorthodox chromosomes’, Current opinion in microbiology, 46, pp. 50–57. https://doi.org/10.1016/j.mib.2018.01.012.

Soyer, J.L. et al. (2020) ‘Genome-wide mapping of histone modifications in two species of Leptosphaeria maculans showing contrasting genomic organization and host specialization’, bioRxiv, p. 2020.05.08.084566. https://doi.org/10.1101/2020.05.08.084566.

Soyer, J.L., Rouxel, T. and Fudal, I. (2015) ‘Chromatin-based control of effector gene expression in plant-associated fungi’, Current Opinion in Plant Biology, 26, pp. 51–56. https://doi.org/10.1016/j.pbi.2015.05.025.

Struhl, K. and Segal, E. (2013) ‘Determinants of nucleosome positioning’, Nature Structural & Molecular Biology, 20(3), pp. 267–273. https://doi.org/10.1038/nsmb.2506.

Szerlong, H.J. and Hansen, J.C. (2011) ‘Nucleosome distribution and linker DNA: connecting nuclear function to dynamic chromatin structure’, Biochemistry and Cell Biology, 89(1), pp. 24–34. https://doi.org/10.1139/O10-139.

Taylor, T.N. and Osborn, J.M. (1996) ‘The importance of fungi in shaping the paleoecosystem’, Review of Palaeobotany and Palynology, 90(3), pp. 249–262. https://doi.org/10.1016/0034-6667(95)00086-0.

Tillo, D. et al. (2010) ‘High nucleosome occupancy is encoded at human regulatory sequences’, PloS One, 5(2), p. e9129. https://doi.org/10.1371/journal.pone.0009129.

Tirosh, I. and Barkai, N. (2008) ‘Two strategies for gene regulation by promoter nucleosomes’, Genome Research, 18(7), pp. 1084–1091. https://doi.org/10.1101/gr.076059.108.

Toruño, T.Y., Stergiopoulos, I. and Coaker, G. (2016) ‘Plant-Pathogen Effectors: Cellular Probes Interfering with Plant Defenses in Spatial and Temporal Manners’, Annual review of phytopathology, 54, pp. 419– 441. https://doi.org/10.1146/annurev-phyto-080615-100204.

Tsankov, A.M. et al. (2010) ‘The role of nucleosome positioning in the evolution of gene regulation’, PLoS biology, 8(7), p. e1000414. https://doi.org/10.1371/journal.pbio.1000414.

Valouev, A. et al. (2008) ‘A high-resolution, nucleosome position map of C. elegans reveals a lack of universal sequence-dictated positioning’, Genome Research, 18(7), pp. 1051–1063. https://doi.org/10.1101/gr.076463.108.

Valouev, A. et al. (2011) ‘Determinants of nucleosome organization in primary human cells’, Nature, 474(7352), pp. 516–520. https://doi.org/10.1038/nature10002.

Van Kan, J.A.L. et al. (2017) ‘A gapless genome sequence of the fungus Botrytis cinerea’, Molecular Plant Pathology, 18(1), pp. 75–89. https://doi.org/10.1111/mpp.12384.

Venters, B.J. and Pugh, B.F. (2009) ‘A canonical promoter organization of the transcription machinery and its regulators in the Saccharomyces genome’, Genome Research, 19(3), pp. 360–371. https://doi.org/10.1101/gr.084970.108.

Wagner, F.R. et al. (2020) ‘Structure of SWI/SNF chromatin remodeller RSC bound to a nucleosome’, Nature, 579(7799), pp. 448–451. https://doi.org/10.1038/s41586-020-2088-0.

Waterhouse, R.M. et al. (2018) ‘BUSCO Applications from Quality Assessments to Gene Prediction and Phylogenomics’, Molecular Biology and Evolution, 35(3), pp. 543–548. https://doi.org/10.1093/molbev/msx319.

Weiberg, A. et al. (2013) ‘Fungal small RNAs suppress plant immunity by hijacking host RNA interference pathways’, Science (New York, N.Y.), 342(6154), pp. 118–123. https://doi.org/10.1126/science.1239705.

Yague-Sanz, C. et al. (2017) ‘A conserved role of the RSC chromatin remodeler in the establishment of nucleosome-depleted regions’, Current Genetics, 63(2), pp. 187–193. https://doi.org/10.1007/s00294-016-0642-y.

Yuan, G.-C. et al. (2005) ‘Genome-scale identification of nucleosome positions in S. cerevisiae’, Science (New York, N.Y.), 309(5734), pp. 626–630. https://doi.org/10.1126/science.1112178.

Zhang, T., Zhang, W. and Jiang, J. (2015) ‘Genome-Wide Nucleosome Occupancy and Positioning and Their Impact on Gene Expression and Evolution in Plants’, Plant Physiology, 168(4), pp. 1406–1416. https://doi.org/10.1104/pp.15.00125.

Zhao, Jing et al. (2021) ‘Distinct Transcriptomic Reprogramming in the Wheat Stripe Rust Fungus During the Initial Infection of Wheat and Barberry’, Molecular plant-microbe interactions: MPMI, 34(2), pp. 198–209. https://doi.org/10.1094/MPMI-08-20-0244-R.

Zhu, P. and Li, G. (2016) ‘Structural insights of nucleosome and the 30-nm chromatin fiber’, Current Opinion in Structural Biology, 36, pp. 106–115. https://doi.org/10.1016/j.sbi.2016.01.013.

## Additional references

Diolez, A. et al. (1995) ‘Boty, a long-terminal-repeat retroelement in the phytopathogenic fungus Botrytis cinerea’, Applied and Environmental Microbiology, 61(1), pp. 103–108. https://doi.org/10.1128/AEM.61.1.103-108.1995.

